# Mitochondrial presequences harbor variable strengths to maintain organellar function

**DOI:** 10.1101/2025.05.26.655807

**Authors:** Youmian Yan, Baigalmaa Erdenepurev, Ian Collinson, Natalie M. Niemi

**Affiliations:** Department of Biochemistry & Molecular Biophysics, Washington University School of Medicine, St. Louis, MO 63110, USA; School of Biochemistry, University of Bristol, Bristol BS8 1TD, United Kingdom

## Abstract

Hundreds of mitochondrial-destined proteins rely on N-terminal presequences for organellar targeting and import. While generally described as positively charged amphipathic helices, presequences lack a consensus motif and thus likely promote the import of proteins into mitochondria with variable efficiencies. Indeed, the concept of presequence “strength” critically underlies biological models such as stress sensing, yet a quantitative analysis of what dictates “strong” versus “weak” presequences is lacking. Furthermore, the extent to which presequence strength affects mitochondrial function and cellular fitness remains unclear. Here, we capitalize on the high-throughput and kinetic nature of the MitoLuc mitochondrial protein import assay to quantify multiple aspects of presequence strength. We find that select presequences, including those that regulate the mitochondrial unfolded protein response (UPR^mt^), are sufficient to impart differential import efficiencies during mitochondrial uncoupling. Surprisingly, we find that presequences beyond those classically associated with stress signaling promote highly variable import efficiency in stressed and basal (i.e., non-stressed) conditions in vitro, suggesting that presequence strength may influence a broader array of processes than currently appreciated. We exploit this variability to demonstrate that only presequences that promote robust import in vitro can fully rescue defects in respiratory growth in Complex IV-deficient yeast, suggesting that presequence strength dictates metabolic potential. Collectively, our findings demonstrate that presequence strength can describe numerous metrics, such as total imported protein, maximal import velocity, or sensitivity to uncoupling, suggesting that the annotation of presequences as “weak” versus “strong” requires more nuanced characterization than is typically performed. Importantly, we find that such variability in presequence strength meaningfully affects cellular fitness in processes beyond stress signaling, suggesting that organisms may broadly exploit presequence strength to fine-tune mitochondrial import and thus organellar homeostasis.

## Introduction

Mitochondria are complex organelles composed of ∼1,000 proteins in yeast^1,2^ and up to 1,500 proteins in mammalian cells and tissues^3–6^. In *Saccharomyces cerevisiae*, all but eight mitochondrial-localized proteins are synthesized outside of the organelle^7^, requiring the targeting and import of hundreds of distinct proteins across the outer mitochondrial membrane. Up to 70% mitochondrial-resident proteins facilitate this localization through an N-terminal mitochondrial targeting signal, or presequence^8,9^. Presequences share common features: they are typically ∼15-50 amino acids in length and enriched in positively charged residues with a tendency to form amphipathic alpha helices^10^. Despite these generally shared traits, presequences are poorly conserved and lack a common sequence motif, rendering it unlikely that they promote protein import with identical kinetics and efficiencies.

Consistently, the concept of presequence strength has been proposed in the literature, with some presequences suggested to be “strong” and others annotated as “weak”^11,12^. Accumulating evidence has highlighted the importance of weak presequences in mitochondrial regulatory strategies^11,13^, including tissue-specific metabolic specialization^14^, mitophagy sensing^15^, and stress responsiveness^16,17^. In these contexts, weak presequences fail to efficiently import cargo proteins into mitochondria, allowing tissue- or context-specific partitioning of a population of protein outside of the organelle to promote specialized mitochondrial regulation. Perhaps the most well-characterized example of such a mechanism underlies the induction of the mitochondrial unfolded protein response, or UPR^mt^, in *Caenorhabditis elegans*. The UPR^mt^ relies on a dual-localized protein, ATFS-1, that contains both a presequence as well as a nuclear localization signal^12^. In normal conditions, ATFS-1 is imported into mitochondria and subsequently degraded by the mitochondrial protease LonP^16^. However, upon mitochondrial stress, the weak presequence of ATFS-1 fails to promote efficient import, allowing an unimported population to translocate to the nucleus and transcriptionally upregulate proteins to mitigate mitochondrial stress, such as the mitochondrial chaperone HSP-60^16–18^. Critically, for the UPR^mt^ to function as proposed, HSP-60 and other mitochondrial-localized stress resolution proteins must harbor strong presequences to enable their translocation into mitochondria, as stressed mitochondria remain compromised for protein import^17,19^. While previous studies have verified the weak import behavior of ATFS-1 in vitro^16,20,21^ and in vivo^22^, the effects of perturbing strong presequences, or any beyond the well-characterized weak ones, are relatively unexplored. Thus, while the differential import behaviors imparted by weak and strong presequences likely critically underlie select stress sensing models, whether mitochondrial presequence strength broadly influences organellar function remains an open question.

One pertinent challenge in understanding the extent to which presequence strength affects mitochondrial function is the lack of a consensus definition of “strong” versus “weak” import behavior. Do strong presequences import proteins at a faster rate than weak presequences? Do they facilitate higher total fractional import, or have higher tolerance to perturbations of mitochondrial membrane potential? Or, do strong presequences mediate more complex behaviors, such as chaperone association or the efficiency of interacting with the import machinery^23,24^? Seemingly, each of these definitions could describe strength, and it is likely that individual presequences display distinct and unique combinations of these metrics. However, many current annotations of presequence strength lack such a nuanced definition, and are rather suggested to be “weak” or “strong” based on their scoring in presequence prediction bioinformatic algorithms such as MitoProtII or MitoFates. The likely complexity of presequence strength suggests that a more rigorous, quantitative approach to define the parameters underlying presequence strength, as well as the identification of contexts in which this behavior is biologically meaningful, is warranted.

One straightforward route for testing presequence strength could involve generating a series of proteins harboring variable presequences fused to a generic cargo protein and comparing their relative import efficiency. While methodologies for quantifying mitochondrial protein import kinetics were developed almost fifty years ago^25,26^, these assays are typically gel-based and rely on autoradiography or western blot-based quantification at a handful of timepoints. Thus, while effective at distinguishing large differences in import efficiencies, these methods lack the resolution and dynamic range to identify more subtle changes in import behavior, and they are limited by the fact that they are low-throughput and difficult to multiplex. Recently, however, a split luciferase-based import assay, MitoLuc, was developed that allows continuous, real-time quantification of in vitro mitochondrial import over a 15- to 30-minute timeframe^27–29^. Given the substantial increase in kinetic resolution and throughput afforded by this assay, we postulated that MitoLuc-based quantification would allow a thorough analysis of import behavior, thus providing a platform for comprehensive assessment of presequence strength.

Here, we perform such quantification using a series of presequences under both basal and uncoupling conditions. We find that presequences are sufficient to promote highly variable import behavior in the MitoLuc assay, which can be quantified by multiple metrics, including total protein imported, maximal import rate, the time required to initiate active import (i.e., the import lag time), and the duration of active import. Using these metrics, we confirm that the presequence of ATFS-1 imparts marked sensitivity to mitochondrial uncoupling, consistent with its proposed role in the UPR^mt^. To test this, we generate a new assay, PotLuc, by multiplexing the <u>pot</u>entiometric dye DiSC_3_(5) with Mito<u>Luc,</u> to find that multiple presequences sustain import during substantial depolarization, suggesting that more presequences than currently appreciated may promote protein translocation in conditions of compromised membrane potential. Beyond these variable responses to uncoupling, we find that seven presequences predicted to be “strong” through bioinformatic algorithms impart highly variable import behavior, displaying differences across all quantified metrics. Through analysis of these metrics of import kinetics, we identify intrinsic features (e.g., presequence charge) and experimentally determined parameters (e.g., precursor thermal stability) that correlate with import behavior in vitro. Finally, we demonstrate that presequences impart differential rescue capacity in yeast with compromised mitochondrial metabolism. Collectively, our study suggests that presequences harbor inherent and complex differences in strength that can be captured in a quantitative manner using the MitoLuc and PotLuc import assays, and that such variability meaningfully affects mitochondrial metabolism and thus cellular fitness.

## Results

### The MitoLuc assay allows high-throughput and quantitative analysis of mitochondrial protein import

To begin, we evaluated the potential of the MitoLuc assay to quantify presequence strength. As mature protein domains can affect mitochondrial protein import efficiency^30^, we generated constructs containing a common cargo protein, DHFR from *Mus musculus*, tagged with a C-terminal epitope encoding the small portion of NanoBiT luciferase (HiBiT). For initial characterization, we used a presequence-containing N-terminal region of yeast Cytochrome b_2_ (Cyb2_Δ43-65_) commonly used in in vitro import assays^31,32^. After purification of this fusion protein from *E*. *coli*, we incubated Cyb2_Δ43-65_-DHFR-HiBiT with isolated yeast mitochondria expressing the large NanoBiT fragment (LgBiT) (Figure 1A). Upon translocation across the inner mitochondrial membrane, the HiBiT-containing Cyb2_Δ43-65_-DHFR can complement the matrix-localized LgBiT, generating a functional NanoBiT that produces light in the presence of its substrate furazine, which can be quantified on a luminescence-compatible plate reader.

**Figure 1.**
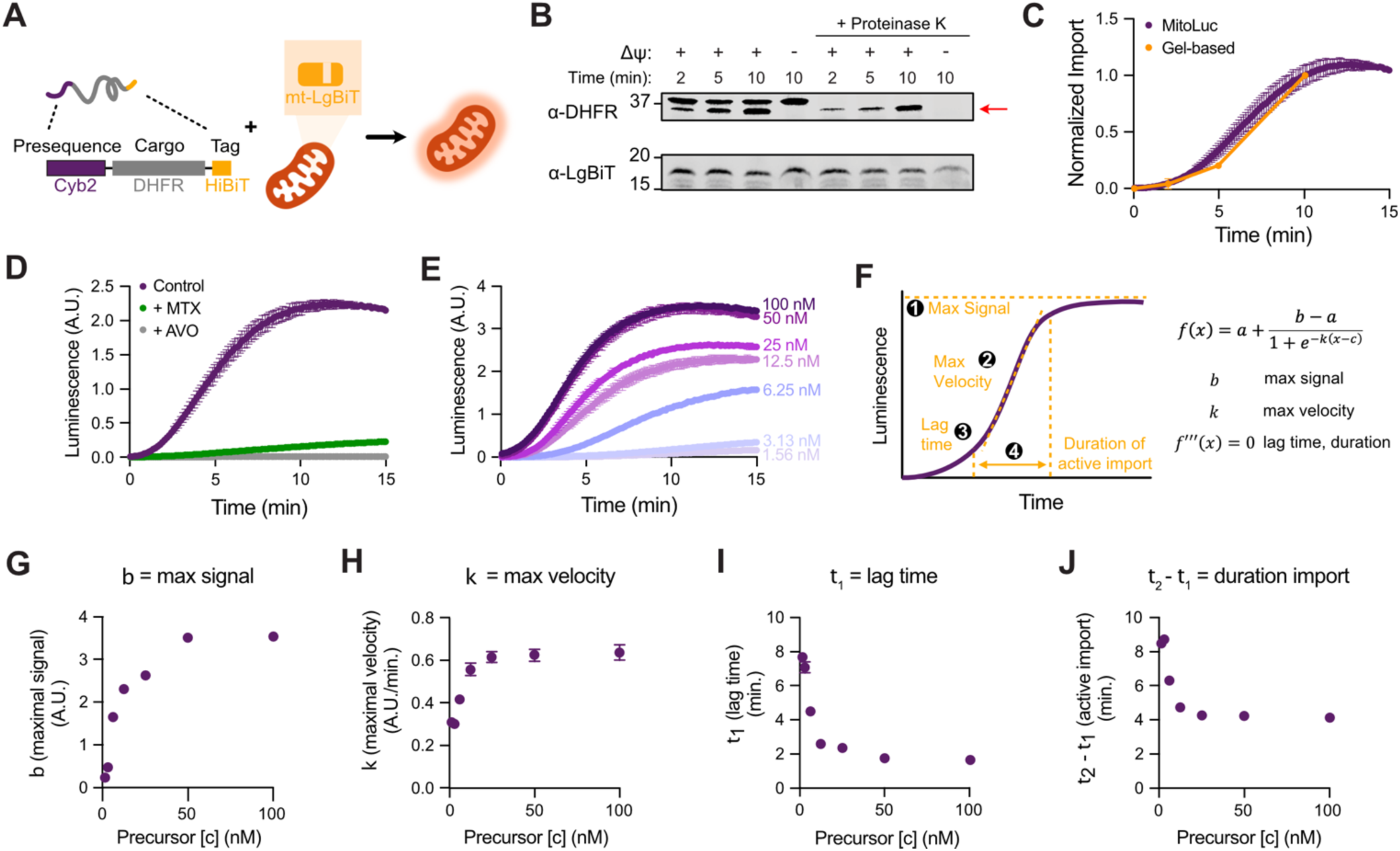
MitoLuc allows quantitative measurements to assess presequence strength. **(A)** Schematic of MitoLuc assay. **(B)** Western blot for the gel-based protein import assay of Cyb2_Δ43-65_-DHFR-HiBiT into isolated mitochondria. Indicated reactions were treated with Proteinase K, and control reactions with dissipated membrane potential (ΔΨ) are shown. **(C)** Quantification of gel-based import (B; 100 nM precursor into 0.2 mg/ml mitochondria) and the MitoLuc assay (12.5 nM precursor into 0.05 mg/mL mitochondria). Fraction of import calculated by normalizing import signal at 10 min. n = 3 independent measurements; data represented as mean ± standard deviation. **(D)** MitoLuc import of 12.5 nM of Cyb2_Δ43-65_-DHFR-HiBiT in untreated (purple), methotrexate (MTX) treated (green) or antimycin A, valinomycin, and oligomycin (AVO, grey) treated conditions. n = 3 independent measurements. Data represented as mean ± standard deviation. **(E)** MitoLuc measurement for the titration of Cyb2_Δ43-65_-DHFR-HiBiT at concentrations ranging from 100 nM to 1.56 nM. n = 3 independent measurements. Data represented as mean ± standard deviation. **(F)** Quantified parameters from the MitoLuc assay with a general logistic function. **(G–J)** Maximal signal *b* (G), maximal velocity *k* (H), lag time *t*_1_ (I) and active duration of import *t*_2_-*t*_1_ (J) for precursor concentrations shown in (E). Each individual curve was fit with a general logistic function and values plotted by precursor concentration. n = 3 independent measurements. Data represented as mean ± standard deviation.

To confirm that the HiBiT tag did not interfere with protein import and that MitoLuc accurately measured mitochondrial protein import, we performed a classic, gel-based import assay. As expected, we detected a time-dependent accumulation of processed Cyb2_Δ43-65_-DHFR-HiBiT protein that was protease-protected (Figure 1B). In parallel, we performed a MitoLuc assay with the identical construct, which resulted in increased luminescence with similar kinetics as the gel-based assay, albeit at mismatched precursor-to-mitochondria ratios (Figure 1C). When comparisons were made between matched precursor-to-mitochondria ratios, the MitoLuc assay exceeded the gel-based assay, emphasizing its enhanced kinetics and dynamic range (Supplemental Figure 1A). Importantly, the luminescence signal was quenched by two treatments expected to diminish protein import: methotrexate (MTX), which binds the DHFR domain to prohibit its unfolding^33^, or a cocktail of antimycin a, valinomycin, and oligomycin (AVO)–disruptors of mitochondrial membrane potential (Figure 1D). These data indicate that the MitoLuc assay reflects bona fide protein import with improved kinetic resolution relative to gel-based assays, motivating us to explore metrics by which import might be quantified using this assay.

To comprehensively characterize import of the precursor protein, we performed a titration of the Cyb2_Δ43-65_-DHFR precursor and found that serial dilutions of Cyb2_Δ43-65_-DHFR-HiBiT showed dose-dependent import behavior (Figure 1E). As MitoLuc generates high-resolution import curves, we sought to fit these data to a model to quantify import behavior. Previous work used a two-step irreversible kinetic model to extract rate constants associated with MitoLuc-quantified import^28^, which worked well for data generated from high precursor concentrations but produced poorer fits at lower precursor concentrations (Supplemental Figure 1B). In particular, the two-step irreversible kinetic model fell short in capturing the initial lag phase (which was more pronounced at lower precursor concentrations) and overestimated the plateau seen at the end of import. To capture this lag phase, we fit the Cyb2_Δ43-65_-DHFR-HiBiT-derived import curves to a generalized logistic function:

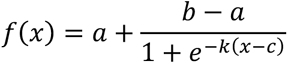

This function contains multiple variables that can quantify precursor import behavior: *b* represents the right maximal asymptote, which reflects the maximal signal of import achieved, while *k* represents the slope of the curve, which quantifies the rate at which import progresses (Figure 1F). To calculate the lag time as well as the total duration of active import, we determined the points of import curves with maximal curvature by solving for the time (*x*) at which the third derivative of the function equals zero (annotated as *t_1_* and *t_2_*, Figure 1F). Whereas the two-step kinetic model captures information about dynamic import behavior that can be used to understand the mechanisms by which import proceeds^28^, the general logistic function provides the best fit to quantify metrics of presequence strength (Supplemental Figure 1C).

Indeed, quantification of import metrics using the general logistic function across precursor concentrations revealed interesting trends. Quantifying maximal import, import rates, lag time, and duration of import for higher Cyb2_Δ43-65_-DHFR-HiBiT concentrations revealed that these parameters approached an asymptote (Figures 1G-J). This suggests that, at higher precursor concentrations, these parameters saturate and the precursor no longer becomes limiting in the MitoLuc import reaction. At lower Cyb2_Δ43-65_-DHFR-HiBiT concentrations, all four parameters showed a dose-dependent response and decreased linearly with precursor concentration (Figures 1G-J). We propose that these parameters can be used to assess presequence strength in a quantitative and rigorous manner.

### The presequences of ATFS-1 and HSP-60 impart similar in vitro basal import behavior but altered sensitivity to mitochondrial stress

Upon establishing quantitative metrics of presequence strength via the MitoLuc assay, we sought to use these parameters to distinguish import behaviors promoted by known “weak” and “strong” presequences. We selected two *C. elegans* proteins involved in the UPR^mt^: ATFS-1 and HSP-60, which have proposed weak and strong presequences, respectively^12^. Though previous studies have confirmed the weak import behavior of ATFS-1 in vitro^16,20,21^ and in vivo^22^, none have directly compared its import behavior to that of *C*. *elegans* stress-responsive proteins such as HSP-60, leaving their relative strengths ill-defined. To test this, we cloned the ATFS-1 or HSP-60 presequence upstream of DHFR-HiBiT (Figure 2A) and quantified the import behavior of these fusion proteins.

**Figure 2.**
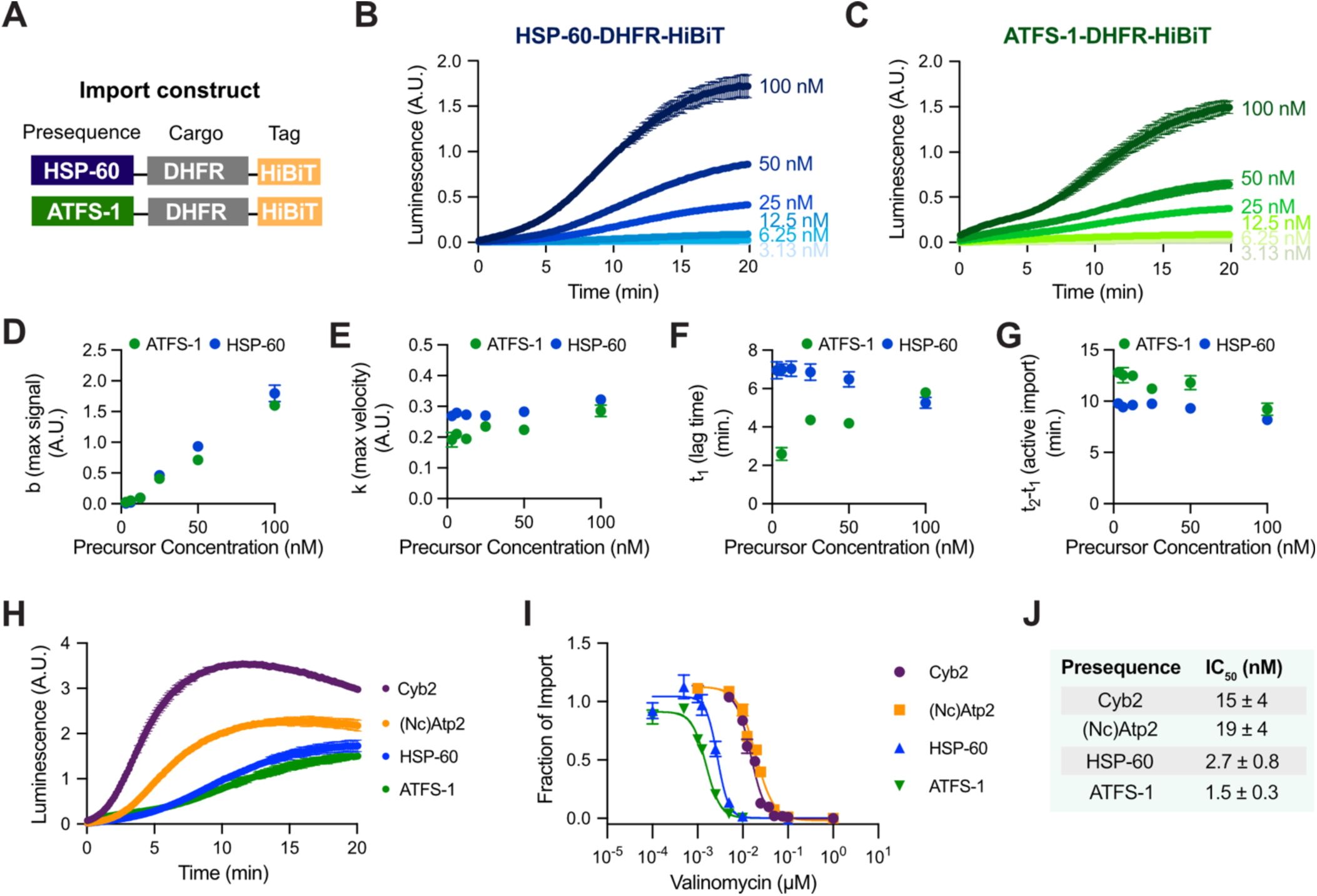
ATFS-1 and HSP-60 presequences promote differential sensitivity to mitochondrial uncoupling. **(A)** Schematic of the HSP-60 and ATFS-1 constructs. **(B, C)** MitoLuc measurement of titrated HSP-60-DHFR-HiBiT (B) and ATFS-1-DHFR-HiBiT (C) precursors from 100 nM to 1.56 nM. n = 3 independent measurements. Data represented as mean ± standard deviation. **(D–G)** Total protein imported (D), maximal velocity (E), lag time (F), and total duration of active import **(G)** across precursor concentrations shown in (B and C). Each parameter reflects individual curves fit with a general logistic function. n = 3 independent measurements. Data represented as mean ± standard deviation. **(H)** MitoLuc import of 100 nM of Cyb2_Δ43-65_-, (Nc)Atp2-, HSP-60-, and ATFS-1-DHFR-HiBiT. n = 3 independent measurements. Data represented as mean ± standard deviation. **(I)** Valinomycin titrations for import of Cyb2_Δ43-65_-, (Nc)Atp2-, HSP-60-, and ATFS-1-DHFR-HiBiT. Data fitted with a four-parameter logistic function. n = 3 independent measurements. Data represented as mean ± standard deviation. **(J)** IC_50_ values for each presequence-containing construct in response to valinomycin.

Surprisingly, we found that titrations of HSP-60-DHFR-HiBiT protein promoted only slightly stronger basal import efficiency than ATFS-1-DHFR-HiBiT (Figures 2B, C). We modeled and fit these import curves using the generalized logistic function at each precursor concentration (Supplemental Figure 2), and found that, unlike Cyb2_Δ43-65_, neither the HSP-60 nor the ATFS-1 presequence showed signs of reaching saturation for any quantified parameter within the concentration range tested (Figures 2D-G). Interestingly, the ATFS-1 presequence gave rise to slower maximal velocity (Figures 2E), which was partially compensated by its shorter lag time (Figure 2F) and longer duration of active import (Figure 2G), leading to only small differences in maximal signals by the end of the import reaction (Figures 2D). To confirm their relatively weak import behavior, we compared both UPR^mt^-associated presequences to Cyb2_Δ43-65_ and the well-characterized presequence of (Nc)Atp2, the beta subunit of ATP synthase in *Neurospora crassa*^30,34^. Consistent with our original findings, both HSP-60 and ATFS-1 presequences promoted much lower basal import of the DHFR cargo than the presequences of Cyb2_Δ43-65_ or (Nc)Atp2 (Figure 2H). Given that the mitochondria used in the MitoLuc assay were isolated from *S. cerevisiae*, it is possible that the ATFS-1 and HSP-60 presequences show lower import rates due to species incompatibility. However, even if such species incompatibility exists, these *C*. *elegans*-derived presequences show similar relative basal import behavior, emphasizing the importance of species-matched comparisons in evaluating presequence strength.

As HSP-60 and ATFS-1 mediate the UPR^mt^, and the ATFS-1 presequence is stress-responsive^16^, we considered these two presequences could show more pronounced differences in import behavior under conditions of uncoupling. Consistent with its role in sensing mitochondrial stress, the ATFS-1 presequence displayed pronounced sensitivity to valinomycin, an uncoupler (Figure 2I, J). Importantly, the HSP-60 presequence showed an elevated IC_50_ relative to ATFS-1, yet fell short of both Cyb2_Δ43-65_ and (Nc)Atp2 (Figure 2I, J). These data reveal several interesting points. First, consistent with its role in the UPR^mt^, the HSP-60 presequence facilitates import in uncoupling conditions in which the ATFS-1 presequence shows diminished import, albeit within a relatively narrow range of valinomycin in the MitoLuc assay. Second, these data demonstrate that basal presequence strength is distinct from presequence strength in uncoupling conditions, as some presequences, such as (Nc)Atp2, show intermediate basal import but strong import capacity during uncoupling (Figures 2H-J). Finally, it is surprising that the four constructs tested had IC_50_ values for valinomycin that spanned an order of magnitude, as this reflects a broader range of uncoupler concentration than might be expected to be necessary to disrupt mitochondrial membrane potential. These observations motivated a more rigorous and quantitative analysis of mitochondrial protein import during conditions of uncoupling.

### Presequences differentially dictate import behavior upon alterations in mitochondrial membrane potential

To quantify import dynamics in response to uncoupling, we used the potentiometric dye 3,3’-dipropylthiadicarbocyanine (DiSC_3_(5)) to assess mitochondrial membrane potential (Δψ). DiSC_3_(5) is a lipophilic molecule that accumulates within polarized membranes, and its fluorescence inversely correlates with Δψ. As expected, addition of mitochondria to DiSC_3_(5)-containing buffer caused robust quenching of fluorescence due to high Δψ of these organelles (Figure 3A). Interestingly, treatment with 10 nM or 100 nM valinomycin moderately increased fluorescence signal, consistent with uncoupling. We could further uncouple these states (Figure 3A), suggesting that these nanomolar valinomycin doses promoted partial uncoupling that may influence the import behavior mediated by the ATFS-1, HSP-60, Cyb2_Δ43-65_, and (Nc)Atp2 presequences. However, to our knowledge, no available assay allows concurrent quantification of protein import and membrane potential. To overcome this challenge, we developed an assay we call PotLuc, which combines measurements from the <u>pot</u>entiometric dye DiSC_3_(5) with the Mito<u>Luc</u> import assay using sequential luminescence and fluorescence reads (Figure 3B). We validated PotLuc using the (Nc)Atp2 presequence, finding normal import kinetics in the presence of DiSC_3_(5) and intact Δψ during import that could be diminished by valinomycin (Figure 3C).

**Figure 3.**
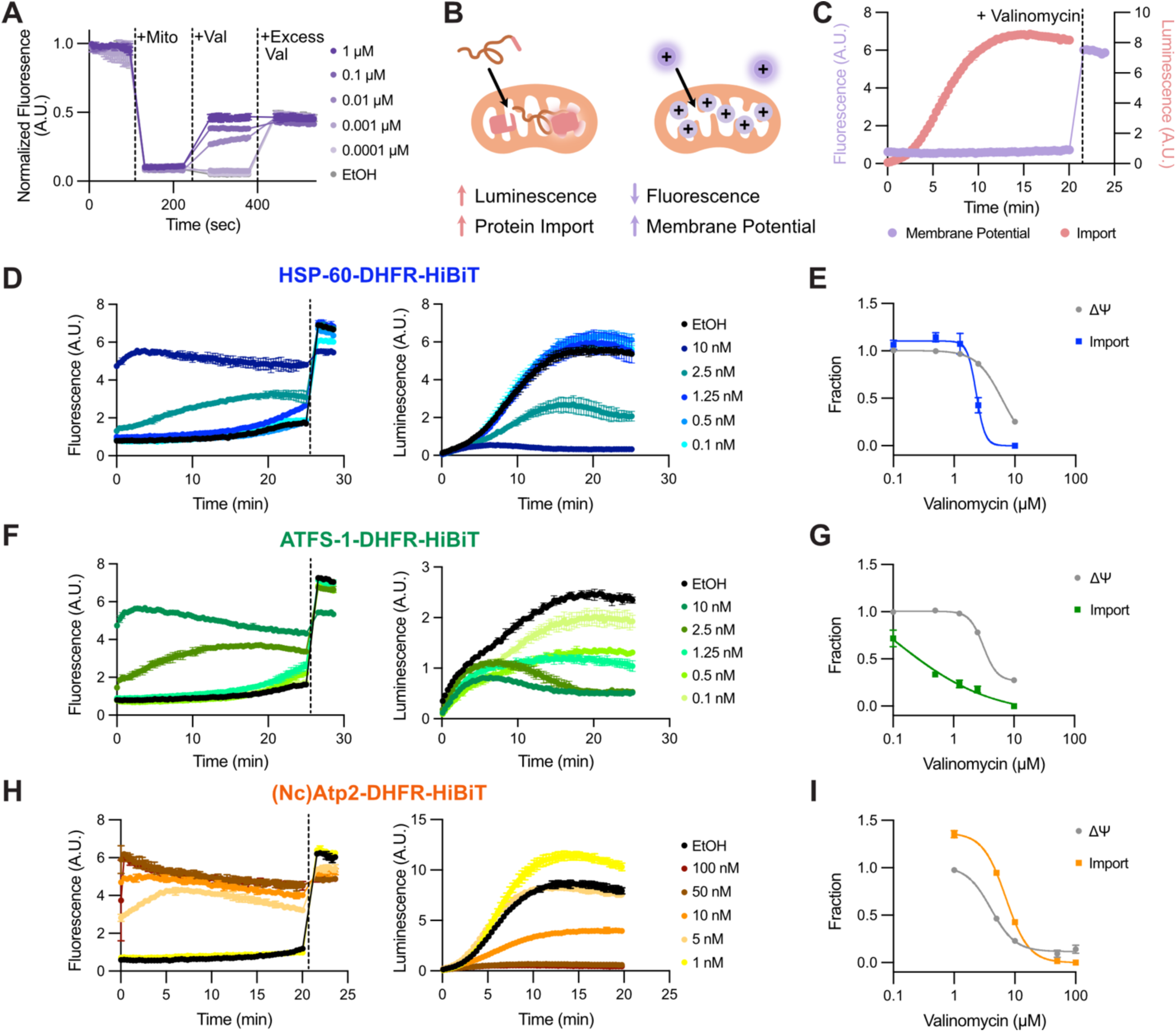
Presequences differentially dictate import behavior upon alterations in mitochondrial membrane potential. **(A)** DiSC_3_(5) fluorescence measurements across valinomycin concentrations. DiSC_3_(5), mitochondria, variable valinomycin, and excess (2 μM) valinomycin sequentially added. Fluorescence was normalized to the maximal signal for each trace. n = 3 independent measurements. Data represented as mean ± standard deviation. **(B)** Schematic of the PotLuc assay. **(C)** Simultaneous quantification of protein import and membrane potential for (Nc)Atp2- DHFR-HiBiT using PotLuc. n = 3 independent measurements. Data represented as mean ± standard deviation. **(D-I)** PotLuc import of 100 nM HSP-60-DHFR-HiBiT (D, E), ATFS-1-DHFR-HiBiT (F, G), and (Nc)Atp2-DHFR-HiBiT (H, I) across valinomycin concentrations and quantifications. Dotted line indicates the addition of 1 μM valinomycin. Data were fitted with a four-parameter logistic function. n = 3 independent measurements. Data represented as mean ± standard deviation.

Given its initial success, we quantified the import behavior of the ATFS-1 and HSP-60 presequences and Δψ during uncoupling using PotLuc (Figures 3D-G). Low valinomycin concentrations had little effect on DiSC_3_(5) fluorescence or the import of HSP-60-DHFR (Figures 3D, E). In contrast, the ATFS-1 presequence showed compromised import in response to even 0.1 nM valinomycin, which notably did not affect DiSC_3_(5) fluorescence (Figures 3F, G). Furthermore, the ATFS-1 presequence showed diminished import capacity even in vehicle only conditions (Figures 3F); as the only difference between the PotLuc and MitoLuc assays is the presence of DiSC_3_(5), this likely reflects mild DiSC_3_(5)-mediated toxicity to which ATFS-1 is sensitive. Collectively, these results indicate that the presequence of ATFS-1 is significantly more sensitive to mitochondrial depolarization than the presequence of HSP-60, consistent with the UPR^mt^ model in which ATFS-1 acts as a sensor for mitochondrial stress and HSP-60 acts as a responder.

Beyond revealing the pronounced sensitivity of the ATFS-1 presequence to uncoupling, we were surprised to see sustained protein import of HSP-60-DHFR-HiBiT in conditions of substantial uncoupling (Figure 3D, teal line). While these data are consistent with the role of HSP-60 as a stress responder, our data indicated that (Nc)Atp2-DHFR-HiBiT and Cyb2_Δ43-65_-DHFR-HiBiT had substantially higher IC_50_ values for valinomycin than HSP-60-DHFR, suggesting these presequences may also allow efficient protein import under compromised Δψ. To test this, we repeated the PotLuc assay with (Nc)Atp2-DHFR-HiBiT and found that this presequence promoted efficient protein import – indistinguishable from a vehicle-only control – in the presence of 5 nM valinomycin, which increased DiSC_3_(5) fluorescence by ∼50% (Figures 3H, peach line; Figure 3I). Even at 10 nM valinomycin, which almost completely dissipated Δψ, the import of (Nc)Atp2-DHFR-HiBiT persisted (Figures 3H, orange line; Figure 3I). Similar behavior was found for the Cyb2_Δ43-65_-DHFR-HiBiT precursor, with protein import seen at greatly compromised Δψ (Supplemental Figure 3A). Protein import in conditions of such substantial uncoupling was surprising, as dissipation of Δψ is amongst the most utilized strategies to block mitochondrial protein translocation. We thus repeated membrane potential quantification with a second potentiometric dye, tetramethylrhodamine ethyl ester (TMRE), obtaining similar changes in relative fluorescence upon valinomycin treatment (Supplemental Figures 3B, C) and indicating the fluorescence measurements of each dye likely reveal bona fide uncoupling rather than technical artifacts. These data suggest that presequences maintain protein import in conditions of compromised membrane potential to varied degrees, further emphasizing the variability in presequence-mediated import behavior in vitro.

### Presequences promote variable mitochondrial protein import efficiency under basal conditions

Our data highlight different presequence strengths in the context of mitochondrial depolarization, leading us to test whether such variation in import kinetics mediated by presequences could manifest in basal (i.e., unstressed) conditions. To test this, we cloned and purified [presequence]-DHFR-HiBiT constructs carrying seven distinct presequences from *S. cerevisiae* proteins (i.e., Hsp60, (Sc)Atp2, Pim1, Mrp21, Mdh1, Cit1, and Mrpl36). Each presequence was selected due to its high MitoFates score, relatively standard length, and sequence confirmation from a study mapping the mitochondrial N-proteome^8^. To fully characterize the import behavior dictated by each presequences, we performed MitoLuc import assays across protein concentrations ranging almost two orders of magnitude (100 nM to 1.56 nM). Consistent with our data on Cyb2_Δ43-65_, HSP-60, and ATFS-1, dose-dependent import behavior was observed for each precursor protein (Figure 4A). However, substantial variability in import efficiency was seen, demonstrating that presequences can drive distinct mitochondrial protein import kinetics under basal conditions. We fit each import curve with the generalized logistic function (Supplemental Figure 4), which revealed several surprising aspects regarding mitochondrial protein import. First, for presequences leading to faster import, such as Hsp60, (Sc)Atp2, and Mrp21, maximal import (*b*) and import velocity (*k*) approached saturation with increasing precursor concentration (Figures 4B, C), analogous to the behavior seen with Cyb2_Δ43-65_ (Figure 1). However, for presequences promoting less efficient import, such as Mdh1 and Mrpl36, these parameters showed a linear response across tested precursor concentrations (Figures 4B, C), similar to the behaviors of the HSP-60 and ATFS-1 presequences (Figures 2B, C). These data suggest that one general property that can distinguish “strong” and “weak” presequences is their ability to saturate quantification metrics within the MitoLuc assay.

**Figure 4.**
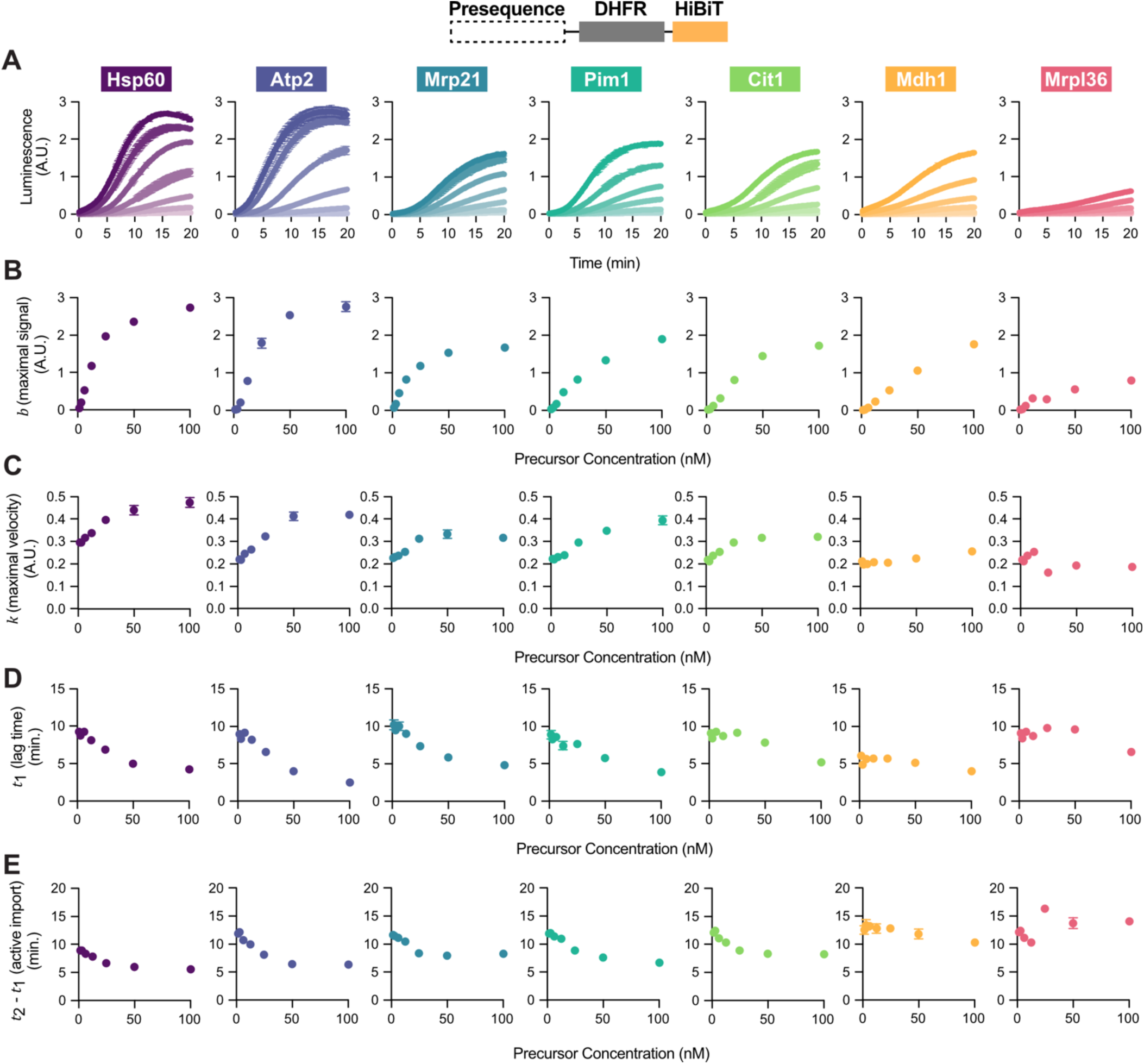
Presequences promote differential protein import efficiency under basal conditions. **(A)** MitoLuc measurements of seven distinct presequence-containing DHFR fusion proteins from *S. cerevisiae* (top). Titration curve of each precursor at 100 nM, 50 nM, 25 nM, 12.5 nM, 6.25 nM, 3.13 nM, and 1.56 nM, with the presequence concentrations varying from high (darkest shade) to low (lightest shade) n = 3 independent measurements. Data represented as mean ± standard deviation. **(B-E)** Parameters extracted from the precursor titration in (A) by fitting individual curves with a general logistic function across precursor concentrations. Results for total protein import (B), maximal import velocity (C), import lag time (D), and duration of active import (E) shown. n = 3 independent measurements. Data represented as mean ± standard deviation.

A deeper analysis of the trends from these MitoLuc quantification metrics indicates that presequence strength can manifest in distinct ways. For instance, though multiple precursors show evidence of saturation for maximal import and velocity, precursors reached distinct saturation plateaus, suggesting presequences carry intrinsic information to influence how fast or how much protein can be imported, at least in vitro. Furthermore, nuanced variabilities in import behaviors across the MitoLuc quantification metrics and across precursor concentrations make it challenging to rank the absolute “strength” of each presequence. As an example, the Mrp21 presequence promoted only intermediate levels of import at 100 nM relative to other precursors (e.g., Hsp60 and (Sc)Atp2), but its import is amongst the fastest at concentrations below 12.5 nM (Figures 4A-C, Supplemental Figure 4). Reciprocally, the Mdh1 presequence also led to intermediate levels of import at 100 nM, but its import capacity declined rapidly with decreasing concentration (Figures 4A-C, Supplemental Figure 4). Precursors with faster import (*k*) tend to have shorter lag times (*t_1_*) that increase with lower protein concentrations (Figure 4D). However, exceptions exist, such as Mdh1, which has short and consistent lag times across tested concentrations. The duration of active import (*t_2_ - t_1_*) showed similar trends to lag time (*t_1_*), with proteins with longer initiation time tending to take longer to complete import (Figure 4E). While the differences in maximal import and maximal velocity are clear among the seven precursors, the differences in lag time and duration of active import are more nuanced, suggesting quantified metrics assess distinct aspects of presequence strength in vitro. Collectively, these data suggest that presequences carry intrinsic information dictating the import capacity of passenger proteins in vitro even under basal conditions, consistent with the concept of presequence strength. However, our data suggest that mitochondrial protein import varies in multiple aspects, such as the amount, rate, and duration of active import, in a concentration- and presequence-dependent manner, and such complexity cannot be fully captured by or reduced to a singular definition of presequence strength.

### MitoLuc quantification identifies predictive metrics for presequence strength

The array of import behaviors displayed across proteins differing only in their presequences raises questions about the molecular determinants giving rise to such variability, and to what extent these determinants may contribute to import behavior. We thus investigated factors previously suggested to affect mitochondrial protein import and determined how well they correlate with each MitoLuc quantification metric (*b*, *k*, *t_1_*, and *t_2_ - t_1_*) derived from 100 nM precursor data fit to the generalized logistic function. A previous study linked protein copy number with mitochondrial protein uptake rates^35^, yet we found little correlation between copy number and MitoLuc import behavior for each of the seven yeast presequences tested (Supplemental Figure 5A). As we purified precursor proteins from *E. coli*, we noted that some preparations contained small but detectable amounts of unknown species matching the molecular weight of bacterial chaperones GroEL and DnaK (Supplemental Figure 5B). As chaperone binding promotes precursor unfolding and thus import^10,24^, we considered that this potential chaperone contamination may influence import rates, yet found that no MitoLuc metrics showed strong correlation with the relative amounts of potential chaperones present (Supplemental Figure 5C).

We next looked to intrinsic properties of presequences to identify factors that may contribute to the variation of basal import quantified by the MitoLuc assay. For each of the ten presequences characterized within our study, we determined parameters previously linked to the efficiency of mitochondrial targeting and import: presequence length, net charge, charge density, and maximal helical hydrophobic moment (μH) (Figure 5A). As presequence strength has been inferred from MitoFates predictions, we also included its scores in our correlation analysis. Notably, MitoFates scores do not correlate with MitoLuc-derived quantification metrics, indicating they are insufficient to describe presequence strength, at least as modeled in the MitoLuc import assays (Figure 5B). Similarly, we find little correlation between presequence length (Supplemental Figure 5D) or charge density (Supplemental Figure 5E) across any MitoLuc quantification metric. While a mild correlation emerged between MitoLuc-based presequence strength and maximal μH (Figure 5C), presequence net charge at pH 7.4 showed the best individual correlation with MitoLuc-derived metrics (Figure 5D), with higher correlations seen for maximal velocity (*k*, R^2^=0.66) and total import duration (*t_2_ - t_1_*, R^2^=0.58) than for total import (*b*, R^2^=0.31) or lag time (*t_1_*, R^2^=0.46). As the total net charge and maximal helical hydrophobic moment could have additive effects in influencing presequence strength, we combined these metrics, finding this combined score correlates to all MitoLuc quantification metrics better than any single tested parameter (Figures 5E, F). Interestingly, the predictive value of this combined metric is higher for the set of weaker presequences (Supplemental Figure 5F). Finally, we determined the stability of each precursor protein through a thermal shift assay (Figure 5G). Though our proteins all contained an identical folded protein domain, we found each presequence-DHFR fusion showed differential thermal stability, albeit in a limited manner relative to methotrexate-induced stabilization (Supplemental Figure 5G). Importantly, these differences in thermal stability inversely correlated with import kinetics, suggesting that relative protein stability likely contributes to, but does not fully explain, import efficiency in the MitoLuc assay. Collectively, these data suggest that both intrinsic presequence properties, such as charge state and maximal helical hydrophobic moment, as well as experimentally determined parameters, such as thermal stability, model presequence strength more accurately than currently used bioinformatic tools.

**Figure 5.**
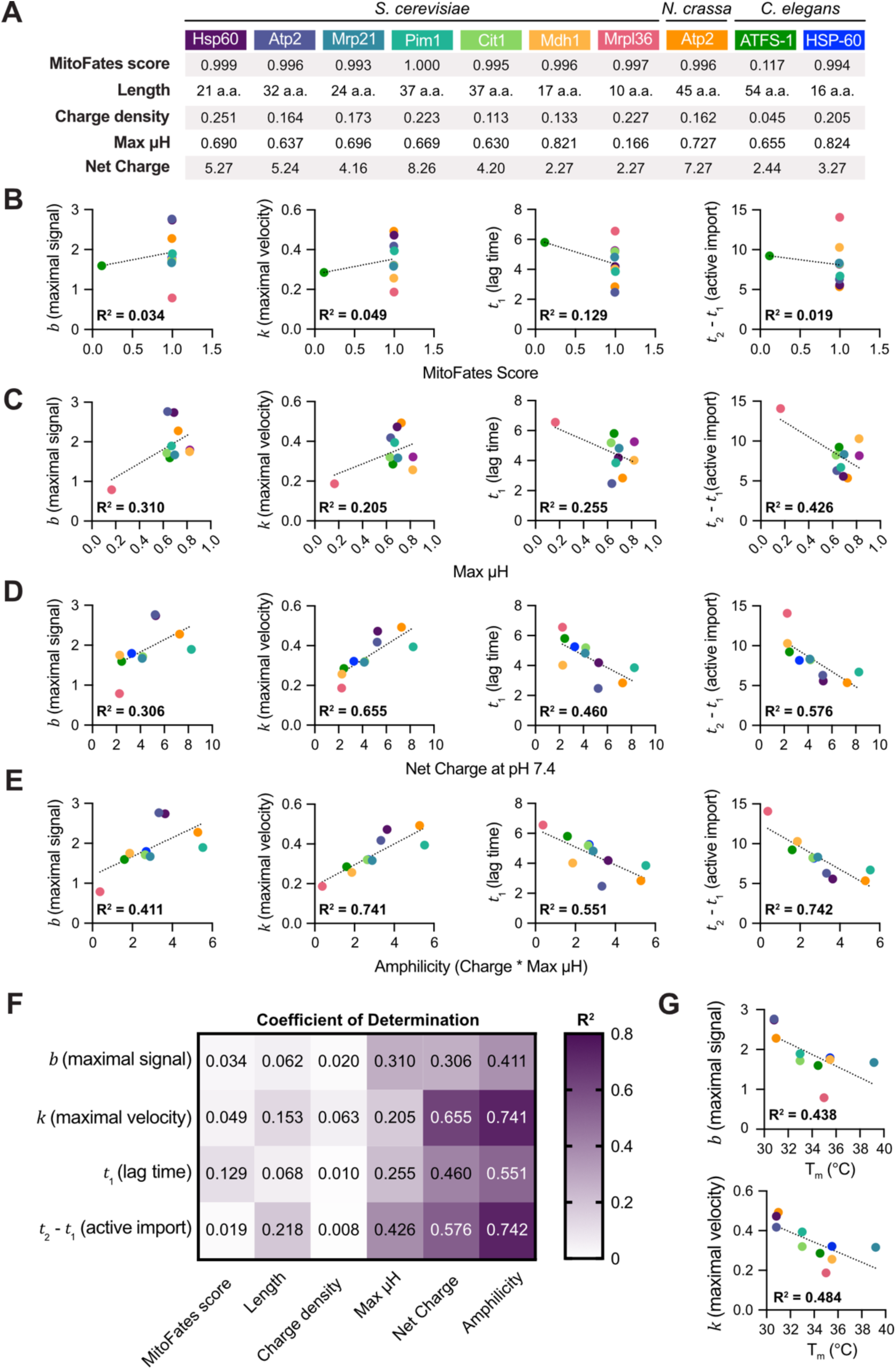
MitoLuc quantification identifies predictive metrics for presequence strength. **(A)** Properties of tested presequences linked to presequence strength. **(B-E)** Correlation between maximal signal (*b*), maximal velocity (*k*), lag time (*t*_1_), or duration of active import (*t*_2_-*t*_1_) and MitoFates scores (B), maximal helical hydrophobic moment (C), net charge at pH 7.4 (D), or the product of maximal helical hydrophobic moment and net charge at pH 7.4 (E). Data fitted with simple linear regression. Color scheme shown as in (A). **(F)** Table showing coefficients of determination for each linear regression comparison. **(G)** Correlation between maximal signal (top) or maximal velocity (bottom) of melting temperatures (T_m_) of the precursor proteins. Data fitted with simple linear regression. Color scheme shown as in (A).

### Strong presequences enable organellar function and organismal fitness

Our results from the MitoLuc and PotLuc assays suggest that intrinsic properties of presequences dictate their ability to promote mitochondrial protein import in vitro, leading us to explore whether such effects could be observed in vivo. To test this, we turned to a classic study that compared presequence-mediated import capacity in *S. cerevisiae* by exploiting variations in growth of *COX4*-null yeast^36^. Cox4 plays a key role in the biogenesis of Complex IV, and, as such, loss of this gene disallows yeast growth on non-fermentable carbon sources. This growth defect can be alleviated by various presequence-Cox4 fusions, including ∼25% of randomly generated sequences tested in the aforementioned study. Importantly, these functional random sequences promoted variable efficiency in rescue, with some mediating “fast” growth and others leading to “slow” growth, indicating a wide range of import-mediated rescue and thus presequence strength. Seeking to exploit this phenotype, we generated *ΔCOX4* W303 yeast, confirmed the mutant yeast could not grow on non-fermentable YPEG plates, and found that this defect could be fully rescued by exogenous full-length Cox4, but not Cox4 lacking a presequence (Figure 6A). One “fast” random presequence selected from the original study rescued growth, but lagged by 2-3 days from wild-type Cox4 rescue (Figure 6A). In comparison, one selected “slow” random presequence showed detectable growth only at days 6-7 of the growth assay (Figure 6A). These data suggest that presequence strength can be inferred by the time required by a presequence-Cox4 fusion protein to rescue *ΔCOX4* yeast growth on a non-fermentable carbon source.

**Figure 6.**
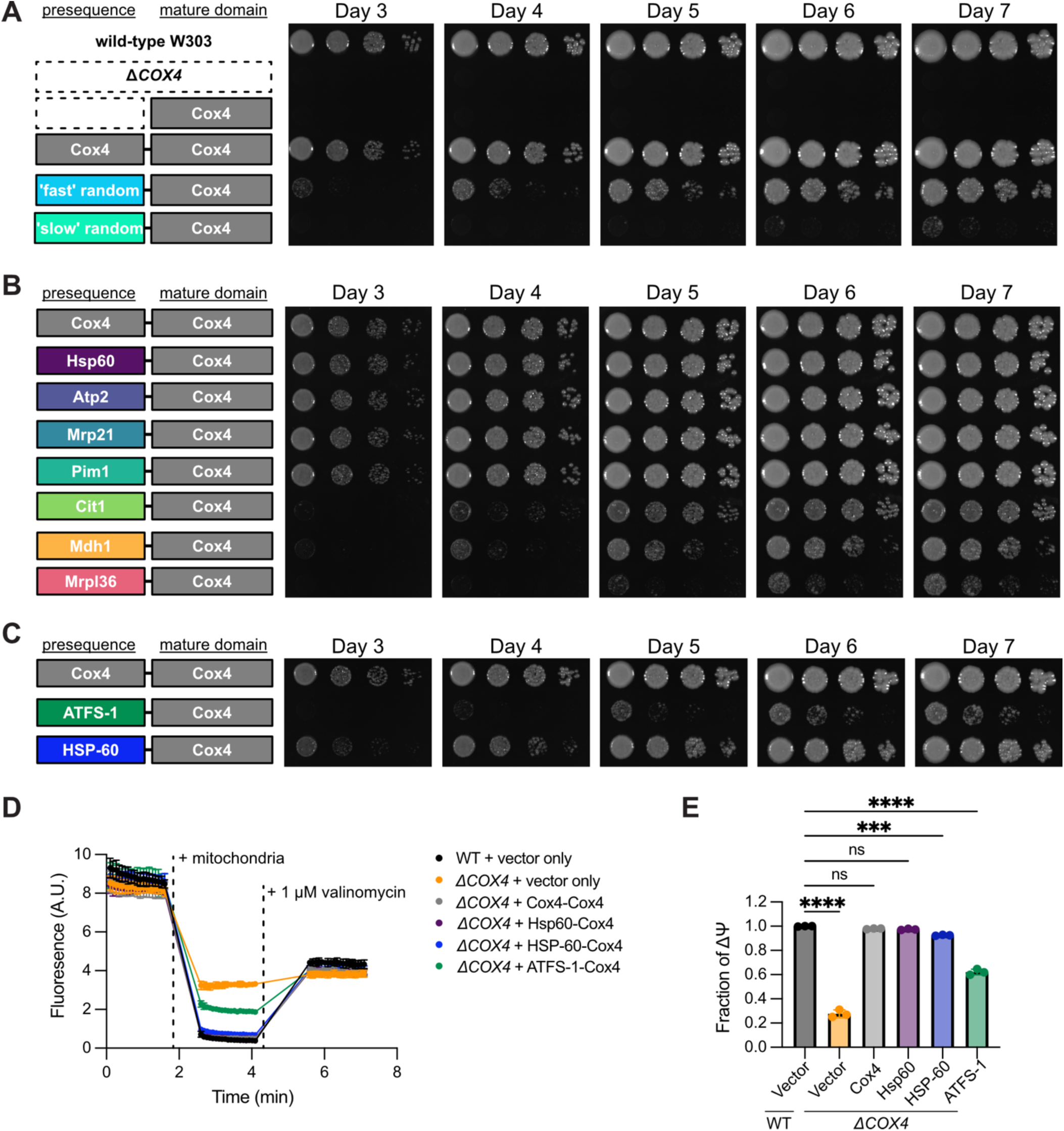
Strong presequences impart full rescue of growth and membrane potential in Δ*COX4* yeast. **(A-C)** Spotting assay of wild-type W303 yeast and *ΔCOX4* yeast rescued with vector or Cox4 fused to various presequences (A), *ΔCOX4* yeast rescued with Cox4 with its endogenous presequence (top) or with the seven tested *S. cerevisiae* presequences (B), or *ΔCOX4* yeast rescued with Cox4 with its endogenous presequence, the ATFS-1 presequence, or the HSP-60 presequence (C). Yeast serial dilutions were plated on YPEG (3% ethanol, 3% glycerol) and incubated at 30°C for the indicated number of days. **(D)** DiSC_3_(5) fluorescence measurement of membrane potential in mitochondria isolated from wild-type (WT), *ΔCOX4*, *ΔCOX4* + Cox4, *ΔCOX4* + (Sc)Hsp60-Cox4, *ΔCOX4* + (Ce)HSP-60-Cox4, and *ΔCOX4* + ATFS-1-Cox4 yeast. DiSC_3_(5), mitochondria, and 1 μM of valinomycin were added as indicated. n = 3 independent measurements. Data represented as mean ± standard deviation. **(E)** Quantification of data in (D). n = 3 independent measurements. Data represented as mean ± standard deviation. ns = not significant, *** p ≤ 0.001, ****p ≤ 0.0001; Ordinary one-way ANOVA.

Given their wide range of basal import kinetics, we hypothesized that the relative strength of each of the seven yeast presequences tested in the MitoLuc assay would proportionately rescue *ΔCOX4* yeast when fused to the Cox4 mature domain. To test this, we replaced the wild-type Cox4 presequence with each of the tested yeast presequences, transformed *ΔCOX4* yeast with each presequence-Cox4 fusion construct, and performed yeast growth assays (Figure 6B). On YPEG plates, presequences of Hsp60, (Sc)Atp2, Mrp21, and Pim1–the four presequences with the strongest import behavior in the MitoLuc assay–rescued yeast growth with the same efficiency as wild-type Cox4 (Figure 6B). The three presequences promoting the weakest import in vitro–Cit1, Mdh1, and Mrpl36–promoted slower growth proportional to their in vitro import capacity. Notably, the weakest tested presequence, Mrpl36, enabled rescue but displayed the slowest growth of all tested constructs, requiring 5 days to promote detectable growth in *ΔCOX4* yeast (Figure 6B)–only slightly better rescue efficiency than the slow random presequence (Figure 6A). Importantly, all strains showed equivalent growth rates on glucose-containing plates, suggesting growth defects seen on non-fermentable carbon sources did not result from impaired viability (Supplemental Figure 6). We also utilized this assay to confirm the strength of the HSP-60 and ATFS-1 presequences; consistent with their relatively slow import kinetics in vitro, neither the ATFS-1 nor the HSP-60 presequence rescued growth as effectively as the wild-type Cox4 presequence (Figure 6C). However, when directly compared, the HSP-60 presequence led to more robust yeast growth than the ATFS-1 presequence (Figure 6C), supporting the UPR^mt^ model in which HSP-60 harbors a “stronger” presequence than ATFS-1.

The difference in rescue mediated by the HSP-60 and ATFS-1 presequences in *ΔCOX4* yeast was surprising, as our initial characterization showed little difference in their abilities to influence basal import capacity (Figure 2). However, we speculated that *ΔCOX4* yeast may harbor defects in mitochondrial membrane potential (Δψ), given the genetic disruption of Complex IV – a proton pumping complex within the electron transport chain. If true, this diminished Δψ could differentially influence HSP-60 and ATFS-1 import efficiency and thus rescue capacity in *ΔCOX4* yeast. To test this, we isolated mitochondria from wild-type yeast, as well as *ΔCOX4* yeast complemented with vector-only, wild-type Cox4, (Sc)Hsp60-Cox4, HSP-60-Cox4, and ATFS-1-Cox4 and measured Δψ with DiSC_3_(5). As predicted, *ΔCOX4* yeast displayed substantially decreased Δψ, which was only moderately rescued by the ATFS-1-Cox4 construct, consistent with its high sensitivity to uncoupling and its slow growth phenotype on YPEG plates (Figures 6D, E). Notably, the strong presequence of (Sc)Hsp60 rescued Δψ to wild-type levels, whereas the *C*. *elegans* HSP-60 presequence imparted high but not full rescue of Δψ (Figures 6D, E), which may explain its growth lag when fused to the Cox4 protein. Overall, the results collected from the *ΔCOX4* rescue assay are consistent with the metrics of presequence strength characterized through MitoLuc and PotLuc assays and generally support the model that presequences encode intrinsic information that dictate the import capacity of their passenger proteins. These data further suggest that the import of Cox4 relies on a strong presequence, which, when substituted for a weak presequence, causes defects in the maintenance of mitochondrial membrane potential and cellular fitness, underscoring the importance of “strong” presequences for maintaining mitochondrial homeostasis under normal physiological conditions.

## Discussion

Up to 70% of the ∼1,000-1,500 mitochondrial-destined proteins carry a presequence to facilitate organellar targeting and import^8,9^. Despite their prevalence, presequences are poorly conserved and have variable compositions and lengths, suggesting many distinct sequence arrangements can target proteins to mitochondria. Indeed, sequences from diverse sources, including the *E*. *coli* genome and random synthetic peptides, are sufficient to promote mitochondrial targeting when fused to a truncated Cox4^36,37^. Furthermore, mapping of the N-terminal mitochondrial proteomes of yeast and mice confirmed that hundreds of cleaved, presequence-containing regions lack a bona fide consensus motif^8,9^. Such divergence across hundreds if not thousands of presequences across evolution raises the interesting and likely possibility that these regions promote mitochondrial import with a wide range of efficiencies.

Indeed, the concept of presequence strength is prevalent in the literature, and multiple “weak” presequences enable the fine tuning of mitochondrial metabolism and responsiveness^14–16^. Most weak presequences have been identified via their low scores in presequence prediction algorithms, such as MitoProtII^38^ or MitoFates^39^. Notably, however, such algorithms were not designed to quantify relative presequence “strength”, but rather to reflect the overall probability that a protein harbors a presequence based on the biochemical properties of its N-terminal region. Through this lens, it is perhaps reasonable to assume that a protein with a low MitoFates or MitoProtII score would harbor a “weak” presequence, as its poor score would suggest its N-terminal region is unlikely to facilitate mitochondrial targeting at all. However, our data suggest that the opposite assumption – that presequences with high MitoFates scores harbor inherently “strong” presequences – is more problematic. We show that seven presequences from *S*. *cerevisiae* with MitoFates scores greater than 0.99 (of a maximal score of 1.0) display remarkable variability in their ability to promote protein import in vitro and in vivo. These data demonstrate that confidence metrics provided by algorithms for the presence of a presequence should not be misinterpreted as proxies for strength and emphasize the importance of in-depth characterization of presequence-mediated import behavior.

One particularly relevant context for quantifying relative presequence strengths lies in the direct comparison of the presequences of ATFS-1 and HSP-60, two proteins whose proposed weak and strong import behavior mediate the mitochondrial unfolded protein response (UPR^mt^) in *C*. *elegans*^12^. While multiple studies have quantified the import kinetics of ATFS-1, each compared its import behavior to the presequence of subunit 9 of the ATP synthase (Su9) of *Neurospora crassa*, one of the strongest presequences characterized in a recent comparative study^24^. Critically, however, the import behavior mediated by the HSP-60 presequence has not been characterized, leaving its strength relative to ATFS-1 unknown. Here, we address this gap in knowledge, revealing that ATFS-1 and HSP-60 have similar import kinetics under basal conditions in the MitoLuc assay. These findings may be biased by potential inefficiencies between *C*. *elegans* presequences and the fungal import machinery, or may reflect bona fide biological behavior independent of species considerations. In either scenario, our data emphasize the importance of making pairwise comparisons most relevant to the biological question of interest. While perhaps initially surprising, the similar basal strengths of ATFS-1 and HSP-60 presequences make sense: in healthy cells, the import of ATFS-1 should not be slower than other mitochondrial proteins, as inefficient import would lead to constitutive activation of the UPR^mt^ even in unstressed conditions. Rather, our data suggest that differences in ATFS-1 and HSP-60 presequence strength emerge in the context of compromised mitochondrial membrane potential, consistent with their differential roles in stress sensing and responsiveness. These data not only signal the need for more thorough analysis to define the relative behaviors of “weak” and “strong” presequences, but also highlight the importance of explicitly defining the behaviors described by presequence “strength”.

The quantitative nature of the MitoLuc assay allowed us to do just this, defining presequence strength in terms of the total amount of protein imported, the maximal rates at which proteins import into mitochondria, the time required to initiate active import, and the duration of active import. Of these, maximal import velocity and the duration of active import correlate well with presequence amphipathicity – a trait long linked to efficient mitochondrial targeting and import^36,40,41^. While many MitoLuc-quantified metrics cluster in “strength” for individual presequences, exceptions exist, and the study of these deviations will likely provide new insights into the nuances of presequence-mediated protein targeting and import. For instance, we find that some presequences that promote intermediate import in basal conditions, such as (Nc)Atp2, can sustain import in conditions of surprisingly strong uncoupling— something similarly seen for the N-terminal region of Cyb2_Δ43-65_. While it is widely accepted that membrane potential is required for mitochondrial protein import, these results raise questions as to the extent of perturbation that can be accommodated to maintain protein import, as well as the magnitude of fluctuation needed to perturb import under physiological conditions. To address these questions, we developed a new assay, PotLuc, which allows simultaneous measurement of protein import with luminescence and membrane potential with fluorescence. Of note, the fluorescence of these potentiometric dyes has been shown to correlate non-linearly with membrane potential, in that a 50% increase in fluorescence may not necessarily correspond to a 50% decrease in absolute membrane potential^42^. Despite this possible limitation, the similar results shown by two independent fluorescent dyes in our study suggest that the compromised membrane potential reported are unlikely to be experimental artifacts, and raise the intriguing possibility that multiple “strong” presequences may sustain import in conditions of compromised membrane potential. Indeed, one might expect that a stress response program would require the import of multiple effectors to mitigate mitochondrial dysfunction. Defining the repertoire and behavior of proteins harboring “strong” stress-responsive presequences will be an important future direction in understanding mitochondrial stress resolution.

Importantly, our data indicate that strong presequences are essential for mitochondrial functions beyond the UPR^mt^. Specifically, we examined *ΔCOX4* yeast as a model of Complex IV deficiency and found that this strain requires a strong presequence to target Cox4 for full rescue. Interestingly, four presequences with the strongest MitoLuc quantified metrics (i.e,. Hsp60, (Sc)Atp2, Mrp21 and Pim1) led to fully restored respiratory growth when fused to a leaderless Cox4 protein, whereas five tested weaker presequences (i.e., Cit1, Mdh1, Mrpl36, HSP-60, and ATFS-1) imbued partial, but not full, rescue of respiratory growth. Mechanistically, this likely derives from a rescue in mitochondrial membrane potential, as Δ*COX4* yeast have compromised Δψ that is fully corrected by the strong presequence of *S*. *cerevisiae* Hsp60, but Δψ is only partially restored by the weak presequence of ATFS-1. The presequence of *C*. *elegans* HSP-60 displayed interesting behavior, promoting a high but not full rescue of Δψ that was insufficient to promote wild-type levels of respiratory growth. This inconsistency highlights the nuance of mitochondrial presequence strength within the complexities of the cellular environment, and may suggest that, in Δ*COX4* yeast, full rescue of membrane potential is required for complete restoration of respiratory growth. Alternatively, these results could suggest that restoration of membrane potential is one of multiple parameters required to maintain cellular fitness in these conditions. It is worth noting that weak tested presequences, such as ATFS-1, still impart respiratory growth in this system, albeit in a lagged fashion. Thus, while weak presequences can promote partial functionality in the rescue of Δ*COX4* yeast, only strong presequences restore cellular fitness to wild-type levels, highlighting the importance of appropriately calibrated presequence strength.

Collectively, our study provides multiple assays, both in vitro and in vivo, to examine presequence strength. Furthermore, we curate our data to demonstrate that a simple calculation using only presequence charge and maximal hydrophobic moment provides a reasonable estimate of maximal import velocity in basal conditions within MitoLuc assay. Such predictions could be applied to virtually any presequence and could establish a more accurate and better resolved metric for presequence “strength” than either the MitoProtII or MitoFates tool, at least for high scoring or “strong” presequences. However, as we have only experimentally characterized a handful of presequences, it is likely that some presequence behavior will deviate from the trends we have observed. We propose that such divergent presequence behavior would be amongst the most important to study, as these exceptions to the rule will likely reveal new complexities to expand our knowledge about the molecular determinants defining presequence strength.

Finally, though the relative strengths of presequences quantified in our in vitro study translate into biologically meaningful behavior when rescuing *ΔCOX4* yeast, it is important to acknowledge the additional complexities of mitochondrial protein import in vivo that cannot be captured by in vitro systems. In the MitoLuc assay, a high concentration of a single protein is loaded into a well-controlled buffering system containing mitochondria devoid of cellular components. Beyond lacking the complexity of the cytoplasm, the in vivo environment would contain copies of multiple presequence-containing proteins, leading to competition for import complex association. Furthermore, bona fide mitochondrial proteins would harbor distinct mature domains with unique properties, such as intrinsic stability or affinity for chaperones, which would influence their targeting, unfolding, and subsequent translocation rates^30,43–45^. Despite these complexities and limitations, we propose that a quantitative understanding of the contributions of presequence strength to import behavior is an important step toward illuminating the regulation of mitochondrial protein import and, ultimately, organellar homeostasis.

## Acknowledgements

This work was supported by the National Institutes of Health (R35GM151130 to N.M. Niemi) and the Wellcome Trust (104632/Z/14/Z to I. Collinson). B. Erdenepurev was supported through the SURGE Summer Undergraduate Research Program within the Department of Biochemistry and Molecular Biophysics at Washington University School of Medicine in St. Louis. We thank Jonathan Friedman (UTSW) and the Niemi Laboratory for careful reading of the manuscript and for helpful discussions on this work. We thank the Pagliarini and Holehouse labs (WUSM) for sharing their plate readers and imagers and the Robertson lab (WUSM) for sharing their sonicator for experiments associated with this work. We thank Junji Suzuki and Yuriy Kirichok (WUSM) for advice and protocols related to the TMRE fluorescence assays. We thank Emily Harrelson (WUSM) for advising on model fitting and for naming the “PotLuc” assay.

## Author contributions

Y.Y and N.M.N. conceived the overall project and its design. Y.Y., B.E., I.C., and N.M.N. generated and validated tools, performed experiments, performed formal analyses, curated data, and/or assisted with data analysis and interpretation of data. Y.Y. and N.M.N. wrote the manuscript and generated the associated figures. Y.Y., B.E., I.C., and N.M.N. reviewed and edited the manuscript.

## Methods

### Supplies and reagents

Methotrexate (catalog #454126), antimycin A (catalog #A8674), valinomycin (catalog #V0627), oligomycin (catalog #O4876), and Prionex Reagent (catalog # 529600) were purchased from MilliporeSigma. Nano-Glo Live Cell Reagent was purchased from Promega (catalog #N2012). Glutathione resin (catalog #NC1057345) and PreScission Protease (catalog #NC1539078) were purchased from Thermo Fisher Scientific. DiSC_3_(5) was purchased from Cayman Chemical (catalog #36308).

### DNA constructs and cloning

pET28-Mff(1-61)-PP-GST was obtained from the laboratory of David Chan (Addgene #73042^46^). pBAD-CyB2_Δ43-65_-HiBiT and pYES-mt-LgBiT were obtained from the laboratory of Ian Collinson^27^. pGEM-SP6-Cyb2_Δ43-65_-DHFR was obtained from the laboratory of Betty Craig. pGEX6P-1-NME2 H118Y was a gift from Stephen Fuhs and Tony Hunter^47^. pJR13019^48^ and pFA6a-His3MX6^49^ were gifts from the laboratory of David Pagliarini. Cyb2_Δ43-65_-DHFR-HiBiT was cloned into the pGEX6P-1 vector, while all other [presequence]-DHFR-HiBiT constructs were cloned into the pET28 vector. [Presequence]-Cox4 constructs were all cloned into the pJR13019 vector with the promoter of Cox4 (i.e., 1 kb upstream of the coding sequence). Plasmids were generated through standard cloning procedures using PCR, restriction enzyme digestion, and ligation. The ligation products were transformed into DH5α competent *E. coli* cells (New England Biolabs, catalog #C2988J), and plasmids were isolated from transformants. All cloned plasmids were validated by Sanger sequencing.

### Yeast cultures

The wild-type haploid W303 (*his3 leu2 lys2 met15 trp1 ura3 ade2*) *Saccharomyces cerevisiae* strain was obtained from the laboratory of David Pagliarini. Δ*COX4* W303 yeast was generated by replacing the COX4 coding sequence with the *his5^+^* selectable marker through a PCR-based method as described previously^49^. Standard lithium acetate-based transformation was carried out for transforming yeast with target plasmids. The spotting assay was carried out on YPEG plate (1% yeast extract, 2% peptone, 3% ethanol, 3% glycerol) or Ura^-^ + 2% glucose plate with 10^4^, 10^3^, 10^2^, and 10^1^ cells plated for each yeast strain. The agar plate was imaged with the ChemiDoc MP Imaging System (Bio-Rad) and Image Lab Touch Software (Bio-Rad, version 3.0.1.14).

### Recombinant protein purification

Recombinant proteins were purified as previously described^29^. The desired expression plasmid was transformed into BL21 (DE3) competent *E. coli* cells (New England Biolabs, catalog #C2527I). Cells were grown in LB media at 37°C on an orbital shaker (225 rpm) until mid-log phase (i.e., OD at 0.6∼0.8), and 1M IPTG was added to induce the protein expression at 18°C for 18 hr (225 rpm). Cells were collected by centrifugation at 4000*g* for 15 min at 4°C. The pellet was resuspended in ice-cold GST lysis buffer (1X phosphate-buffered saline, 1% (v/v) Triton X, 5% (v/v) glycerol, 1 mM dithiothreitol (DTT), 10 mL/500 mL bacterial culture). After lysing the cells through sonication, the mixture was centrifuged at 14,000*g* for 30 min at 4°C, and the supernatant was kept. Glutathione resin (1.5 mL resin/500 mL bacterial culture) was washed with 10X bed volume of GST lysis buffer for three times through pelleting (3000*g* for 1 min) and resuspending at 4°C. The collected supernatant of cell lysates was incubated with the washed resins at 4°C for 2.5 hr with rotation. The resins were washed with 10X bed volume of GST lysis buffer for three times and then with 10X bed volume of PreScission Protease buffer (20 mM Tris-HCl, 150 mM NaCl, 0.5 mM ethylenediaminetetraacetic acid (EDTA), 1 mM DTT, pH 7.4) three times. The resins were resuspended in 1X bed volume of PreScission Protease buffer, and PreScission Protease was added (12.5U/500 mL bacterial culture). The mixtures were incubated at 4°C overnight with rotation. The supernatant was collected by centrifuging at 3000*g* for 5 min at 4°C, and proteins were concentrated using centrifugal concentrators with a molecular weight cut-off of 30K for Cyb2_Δ43-65_-DHFR-HiBiT and 10K for all other [presequence]-DHFR-HiBiT constructs (Vivaproducts, catalog #VS0621, VS0601, VS2001). After aliquoting, the proteins were stored in −80°C until use in import assay. The protein concentration was approximated by running the samples on a 10% SDS-PAGE gel with BSA standards (Thermo Fisher Scientific, catalog #PI23225) and calculating from the standard curve.

### Crude mitochondrial enrichment

Mitochondria were isolated as previously described^29^. For induction of mt-LgBiT, a single colony of yeast transformed with pYES-mt-LgBiT were incubated in 50 mL Ura^-^ + 2% glucose media at 30°C overnight on an orbital shaker (225 rpm). 4×10^8^ cells were diluted into 500 mL YPG media (1% yeast extract, 2% peptone, 3% glycerol), and the yeast were grown at 30°C (225 rpm). Yeast were grown to mid-log phase (i.e., OD at 0.6∼0.8) and 1% galactose was added to induce mt-LgBiT expression for 3 hr before harvesting. For *ΔCOX4* yeast strains and wild-type W303 yeast, a single colony of yeast transformed with desired plasmids were incubated in 50 mL His^-^Ura^-^ + 2% glucose media at 30°C overnight on an orbital shaker (225 rpm). 4×10^8^ cells were diluted into 500 mL YPGal media (1% yeast extract, 2% peptone, 3% galactose), and yeast were grown at 30°C (225 rpm) overnight (for wild-type + vector, *ΔCOX4* + Cox4, *ΔCOX4* + Hsp60-Cox4, *ΔCOX4* + HSP-60-Cox4) or for two days (*ΔCOX4* + vector, *ΔCOX4* + ATFS-1-Cox4). Cells were collected through centrifugation at 3000*g* for 5 min and washed with sterile water. After spinning down the cells (3000*g* for 5 min) and determining the wet weight, the cell pellet was resuspended in DTT buffer (100 mM Tris-H_2_SO_4_, 10 mM DTT, pH 9.4, 2 mL/g wet weight cells) and then shaken slowly (80 rpm) at 30°C for 20 min. After spinning down the yeast at 3000*g* for 5 min, the cell pellet was washed with zymolase buffer (1.2 M sorbitol, 20 mM potassium phosphate, pH 7.4, 7 mL/g wet weight cells). After another spin at 3000*g* for 5 min, the pellet was resuspended in zymolase buffer (7 mL/g wet weight cells) containing zymolase (3 mg/g wet weight cells, Thermo Fisher Scientific, catalog #NC0497252) and then shaken slowly (80 rpm) at 30°C for 45 min. The cells were pelleted (3000*g,* 5 min) and washed with zymolase buffer (7 mL/g wet weight cells). After another spin (3000*g*, 5 min), the cell pellet was resuspended in ice-cold homogenization buffer (0.6 M sorbitol, 10 mM Tris-HCl, 1 mM ethyleneglycol-bis(β-aminoethyl)-N,N,Nʹ,Nʹ-tetraacetic acid (EGTA), 1 mM phenylmethylsulfonyl fluoride, 0.2% (w/v) fatty acid-free bovine serum albumin (BSA), pH 7.4, 6.5 mL/g wet weight cells). The yeast cells were homogenized using a glass Dounce homogenizer with 8 strokes and then diluted twofold with the homogenization buffer on ice. The resulting mixture were centrifuged at 1500*g* for 5 min at 4°C, and the supernatant was kept and centrifuged at 4000*g* for 5 min at 4°C. The supernatant was collected and centrifuged at 12,000*g* for 15 min at 4°C. The resulting mitochondrial pellet was then resuspended in ice-cold SEM buffer (250 mM sucrose, 1 mM EGTA, 10 mM MOPS-KOH, pH 7.2) with wide-bore tips to reach a final protein concentration around 5∼10 mg/mL. The concentration was determined using the Pierce 660 nm Protein Assay Kit (Thermo Fisher Scientific, catalog #22662). After aliquoting, the samples were flash-frozen in liquid nitrogen and stored in −80°C until use in import assay or membrane potential measurement.

### Gel-based mitochondrial import assay

Import assays were performed based on previously published protocol^50^. Mitochondria were resuspended in ice-cold import buffer (250 mM sucrose, 80 mM potassium chloride, 1 mM potassium phosphate, 5 mM magnesium chloride, 10 mM MOPS-KOH, 0.1% (v/v) Prionex reagent, 2 mM NADH, 1 mM ATP, 0.1 mg/mL creatine kinase, 5 mM phosphocreatine, pH 7.2) to reach a final protein concentration of 0.2 mg/mL. Where indicated, 10 μM antimycin A, 1 μM valinomycin, and 20 μM oligomycin (AVO) were added to dissipate the mitochondrial membrane potential. The mixtures were incubated at room temperature for 2 min, and the import reactions were initiated by addition of 0.1 μM of purified CyB2_Δ43-65_-DHFR-HiBiT.

After the specified time, the import reactions were terminated by addition of AVO. Each mixture was then split to two, and 10 μg/mL of Proteinase K was added to half of them to remove non-imported precursor proteins. After incubation on ice for 15 min, mitochondria were collected by centrifugation at 12,000*g* for 5 min at 4°C. The pellet was resuspended in sample buffer (62 mM Tris-HCl pH 6.8, 5% (w/v) sucrose, 1% (w/v) sodium dodecyl sulfate, 0.01% (w/v) bromophenol blue, 1% (v/v) β-mercaptoethanol) and boiled at 95°C for 5 min. The samples were then run on a 15% SDS-PAGE gel at 120V for 90 min with PreScission Plus All-Blue Prestained Protein Standards (Bio-Rad, catalog #1610373). samples were transferred to a nitrocellulose membrane (Thermo Fisher Scientific, catalog #45004016) at 100V for 60 min at 4°C. After blocking with 3% nonfat dairy milk in TBS-T, membranes were incubated with primary antibodies overnight at 4°C. Primary antibodies used are anti-DHFR (Cell Signaling Technology, catalog #43497, dilution 1:1000) and anti-LgBiT (Promega, catalog #N7100, dilution 1:1000 after reconstitution to 0.5 mg/mL). Membranes were washed three times by incubating in TBS-T for 5 min and then incubated with secondary antibodies conjugated with fluorophores (LI-COR Bioscience, catalog #926-32210 and 926-32211) for 30 min at room temperature. Membranes were washed three times by incubating in TBS-T for 5 min and then imaged with a LI-COR OdysseyFC instrument and Image Studio software (LI-COR Bioscience, version 5.2).

#### MitoLuc assay

A detailed protocol for performing MitoLuc import assays has been published previously^29^. Briefly, 2X Import Buffer (500 mM sucrose, 160 mM potassium chloride, 2 mM potassium phosphate, 10 mM magnesium chloride, 20 mM MOPS-KOH, 0.2% (v/v) Prionex reagent, pH 7.2) was prepared and cooled to 4°C. For each import reaction, Mixture 1 which was 1.25X final concentration (62.5 μg/mL of mitochondria in 1X Import Buffer) and Mixture 2 which was 5X final concentration (5X precursor proteins, 1.25X Nano-Glo luciferase assay substrate (supplied at 100X, final concentration 0.25X) in 1X Import Buffer) were prepared on ice. For AVO treatment, 12.5 μM antimycin A, 1.25 μM valinomycin, and 25 μM oligomycin was added to Mixture 1. For methotrexate treatment, 2.5 μM of MTX was added to Mixture 2 and incubated on ice for 5 min. For valinomycin titration, 1.25X of the target concentration was added to Mixture 1. To initiate the import reaction, 100 μL of Mixture 1 was mixed with 25 μL of Mixture 2 on a white flat-bottom 96 well plate (Pro Lab Supply, catalog #30196) at room temperature. Luminescence was measured with a BioTek Synergy LX Multimode Reader (Agilent Technologies) using the luminescence filter and BioTek Gen5 software (Agilent Technologies, version 3.11). Measurements were taken every 8 seconds, and the plate was shaken linearly for 1 second before each measurement. Acquisition time was set to 0.2 second with a gain of 100. For fitting to the general logistic function using non-linear least square analysis, data points after reaching the maximal luminescence signal for each import trace were excluded. For the four parameters, [*a*, *b*, *c*, *k*], constraints were set as follows: upper limit was set to [Infinite, 6, 20, 1], lower limit was set to [-3, 0, 2, 0], and the initial value was set to [-0.01, 1, 10, 0.5]. Model fitting and solving for third derivatives were done in MATLAB (MathWorks, version R2024b), and Prism (Graphpad, version 10.4.2) and Affinity Designer 2 (Serif, version 2.6.3) were used to generate the figures. For valinomycin titration in Figure 2, the import of 12.5 nM of CyB2_Δ43-65_-DHFR-HiBiT, 25 nM of (Nc)Atp2-DHFR-HiBiT, 100 nM of HSP-60-DHFR-HiBiT, or 100 nM of ATFS-1-DHFR-HiBiT was measured at varied concentrations of valinomycin as specified. For fraction of import, the maximal luminescence signal was used to represent the amount of proteins imported (*L_max_*) for each treatment. The maximal signal for the trace at the highest valinomycin concentration for each construct was used to represent the background signal (*L_background_*), and the maximal signal for the trace with EtOH treatment was used as maximal import for normalization (*L_EtOH_*). The fraction of import was then calculated using the equation,

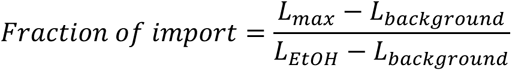

*L_max_* varies based on the valinomycin concentration, while *L_background_* and *L_EtOH_* were constant for all calculations for each precursor protein. Calculations were done in Excel, and Prism was used to generate the figures. The fraction of import data was fitted with the “[Inhibitor] vs. response --Variable slope” model in Prism using non-linear least square regression. The standard deviation was calculated from the 95% confidence interval generated from fitting.

### Membrane potential measurement

Mitochondria were resuspended in ice-cold 1X Import Buffer (250 mM sucrose, 80 mM potassium chloride, 1 mM potassium phosphate, 5 mM magnesium chloride, 10 mM MOPS-KOH, 0.1% (v/v) Prionex reagent, pH 7.2) to a final protein concentration of 0.25 mg/mL. 100 μL of 1X Import Buffer containing 1 μM of DiSC_3_(5) was added to a black-wall clear-bottom 96 well plate (Thermo Fisher Scientific, catalog #07-000-166), and fluorescence was monitored with a BioTek Cytation 5 Cell Imaging Multimode Reader (Agilent Technologies) and BioTek Gen5 software (Agilent Technologies, version 3.15). After linear shaking for 5 seconds, fluorescence was measured for 1.5 min at a 5-second interval with excitation at 622 nm and emission at 670 nm. Then, 25 μL of the resuspended mitochondria were added to reach a final concentration of 0.05 mg/mL. The plate was shaken linearly for 5 seconds, and fluorescence was measured for 1.5 min with the same setting. For titration of valinomycin, varied concentrations of valinomycin were added to reach the specified final concentration, and the plate was shaken linearly for 5 seconds, followed by fluorescence measurement for 1.5 min with the same setting. Lastly, 2 μM of valinomycin was added to completely dissipate the membrane potential. After 5 seconds of linear shaking, fluorescence was measured for 1.5 min with the same setting. For quantification of membrane potential without valinomycin titration, 1 μM of valinomycin was added after fluorescence measurement in the presence of mitochondria to dissipate the membrane potential. To calculate fraction of Δψ shown in Figure 6, the average fluorescence signal over the last 1 min within the 1.5 min measurement window after mitochondria addition was calculated for the effects of mitochondria (*F_mito_*) for each measurement, and the average fluorescence signal over the last 1 min within the 1.5 min measurement window after 1 μM valinomycin addition was calculated only for the mitochondria isolated from wild-type + vector yeast for maximal fluorescence recovery (*F_max_*). Fraction of Δψ was then calculated using the equation,

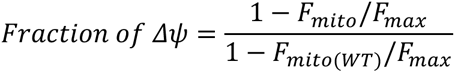

*F_mito_* varies based on the yeast strain from which mitochondria were isolated, and *F_max_* and *F_mito(WT)_* came from the measurement of mitochondria isolated from wild-type + vector yeast for normalization. All calculations were done in Excel, and Prism and Affinity Designer 2 were used to generate the figures. For measurement using TMRE, the procedures were the same with the exceptions as follows: 200 nM of TMRE was used instead of 1 μΜ of DiSC_3_(5), fluorescence was measured with excitation at 549 nm and emission at 574 nm, and each measurement window was elongated to 2 min from 1.5 min.

### PotLuc assay

The procedure and solution composition were the same as the MitoLuc assay with the exceptions listed below. 1.25 μM of DiSC_3_(5) was added to Mixture 1. The samples were loaded to a black-wall clear-bottom 96 well plate and measured with a BioTek Cytation 5 Cell Imaging Multimode Reader and BioTek Gen5 software. Measurements were taken every 20 seconds, and the plate was shaken linearly for 1 second before each measurement. Fluorescence was measured with excitation at 622 nm and emission at 670 nm, and luminescence was measured with acquisition time of 0.2 seconds and a gain of 100. After 25 min (ATFS-1-DHFR-HiBiT and HSP-60-DHFR-HiBiT) or 20 min ((Nc)Atp2-DHFR-HiBiT and Cyb2_Δ43-65_-DHFR-HiBiT) of measurement to let the import reaction complete, 1 μM of valinomycin was added to dissipate the membrane potential. After 5 seconds of linear shaking, fluorescence was measured for 2 minutes at an interval of 20 seconds without shaking. To calculate fraction of Δψ shown in Figure 3, the average fluorescence signal over the first 1 min within the 25 min or 20 min import reaction measurement was calculated for the effects of varied dosage of valinomycin (*F_val_*) for each measurement, and the average fluorescence signal over the first 1 min after addition of 1 μM valinomycin at the end was calculated only for the EtOH control treatment for maximal fluorescence recovery (*F_max_*). Fraction of Δψ was then calculated using the equation,

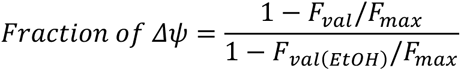

*F_val_* varies based on the valinomycin concentration, and *F_val(EtOH)_* and *F_max_* came from the measurement of EtOH treatment for normalization. Fraction of import was calculated the same way as specified in MitoLuc assay section above. Calculations were done in Excel, and Prism and Affinity Designer 2 were used to generate the figures. For fraction of import and fraction of Δψ, the data were fitted with the “[Inhibitor] vs. response --Variable slope” model in Prism using nonlinear regression.

### Protein thermal shift assay

Recombinant proteins were diluted in PreScission Protease Buffer (20 mM Tris-HCl, 150 mM NaCl, 0.5 mM EDTA, 1 mM DTT, pH 7.4) containing 38 μM of methotrexate or vector (1.25% (v/v) DMSO in PBS) to reach a final protein concentration of 0.2 mg/mL. For Mrp21-DHFR-HiBiT, a final concentration of 0.11 mg/mL was used due to low yield of protein purification. The samples were incubated on ice for 10 min before addition of SYPRO Orange Protein Gel Stain (Thermo Fisher Scientific, catalog #S6650, supplied at 5000X, final concentration 5X). The mixtures were then loaded on a low profile non-skirted 96-well PCR plate (Thermo Fisher Scientific, catalog #AB-0700) with 25 μL of sample per well. The plate was sealed with a qPCR optical grade plate seal (Thermo Fisher Scientific, catalog #AB-1170). Fluorescence was measured with a CFX96 Touch Real-Time PCR Detection system (Bio-Rad) and the CFX Maestro software (Bio-Rad, version 2.3) from 10°C to 95°C in increments of 0.5°C with 10 seconds at each temperature, following the protocol from Bio-Rad. Data was analyzed in Excel, and Prism and Affinity Designer 2 were used to generate the figures.

---

**Supplemental Figure 1.**
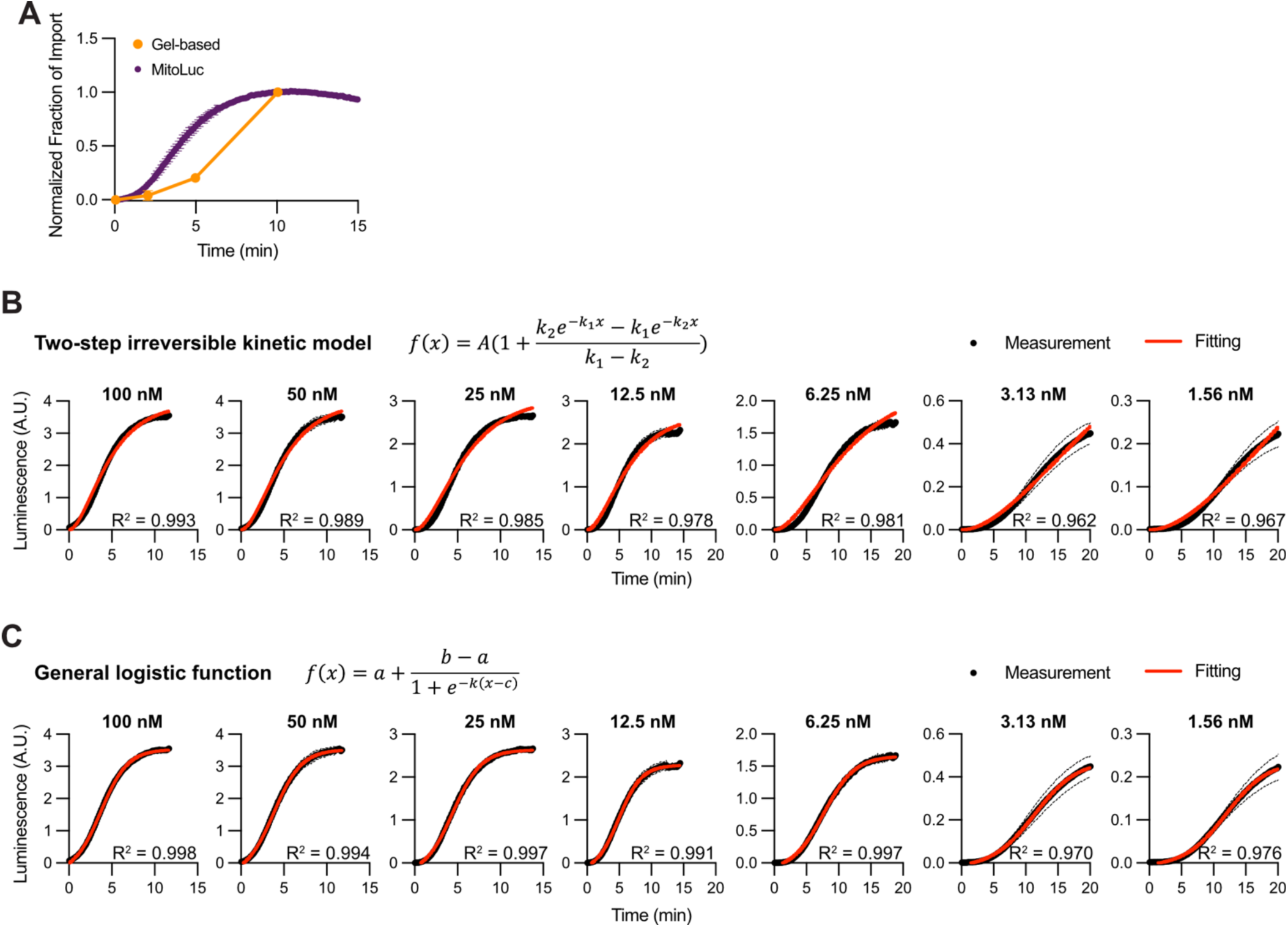
**(A)** Quantification of the gel-based import assay shown in Figure 1 (B) and the MitoLuc assay for the import of 100 nM (gel-based) or 25 nM (MitoLuc) of Cyb2_Δ43-65_-DHFR-HiBiT into 0.2 mg/mL (gel-based) or 0.05 mg/mL (MitoLuc) isolated mitochondria. Fraction of import was calculated by normalizing to the signal at 10 min of import. n = 3 independent measurements. Data represented as mean ± standard deviation. **(B)** Import traces of Cyb2_Δ43-65_-DHFR-HiBiT from the MitoLuc assay were fitted with the two-step irreversible kinetic model at each precursor concentration as reported previously^28^. n = 3 independent measurements. Data represented as mean ± standard deviation. **(C)** Import traces of Cyb2_Δ43-65_-DHFR-HiBiT from the MitoLuc assay were fitted with the generalized logistic function at each precursor concentration. n = 3 independent measurements. Data represented as mean ± standard deviation.

**Supplemental Figure 2.**
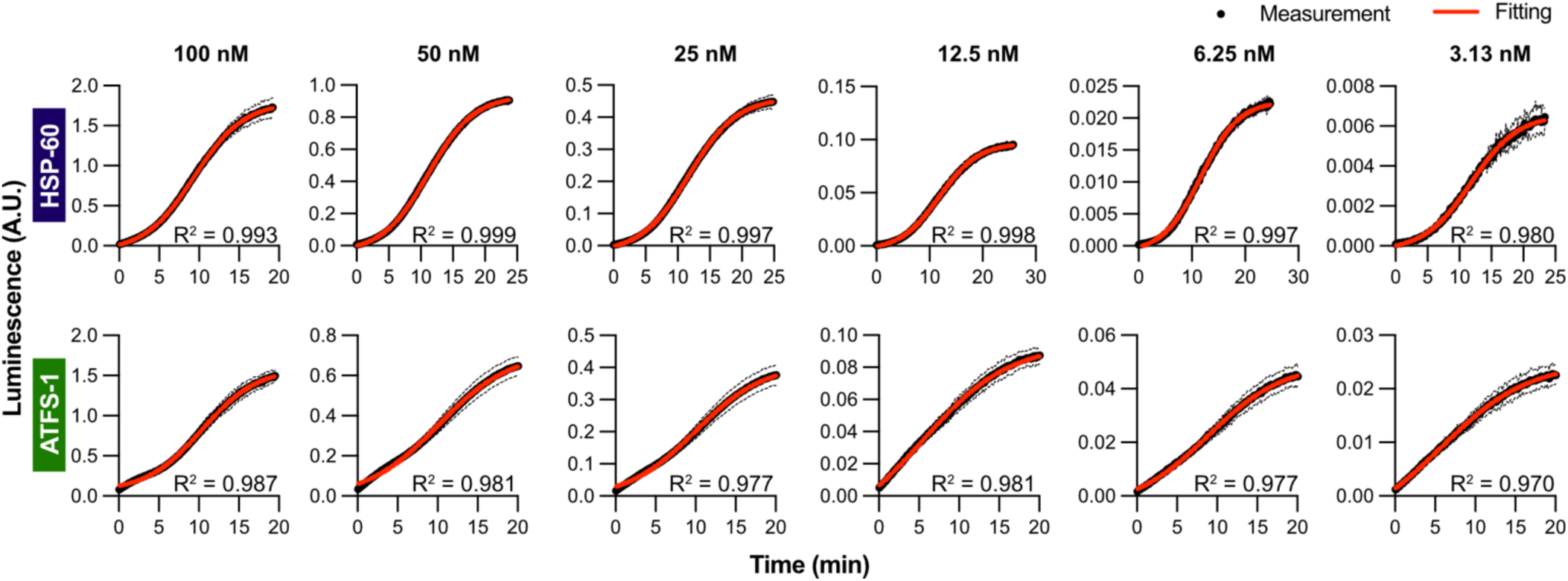
Import traces of HSP-60-DHFR-HiBiT and ATFS-1-DHFR-HiBiT from the MitoLuc assay were fitted with the generalized logistic function at each precursor concentration. n = 3 independent measurements. Data represented as mean ± standard deviation.

**Supplemental Figure 3.**
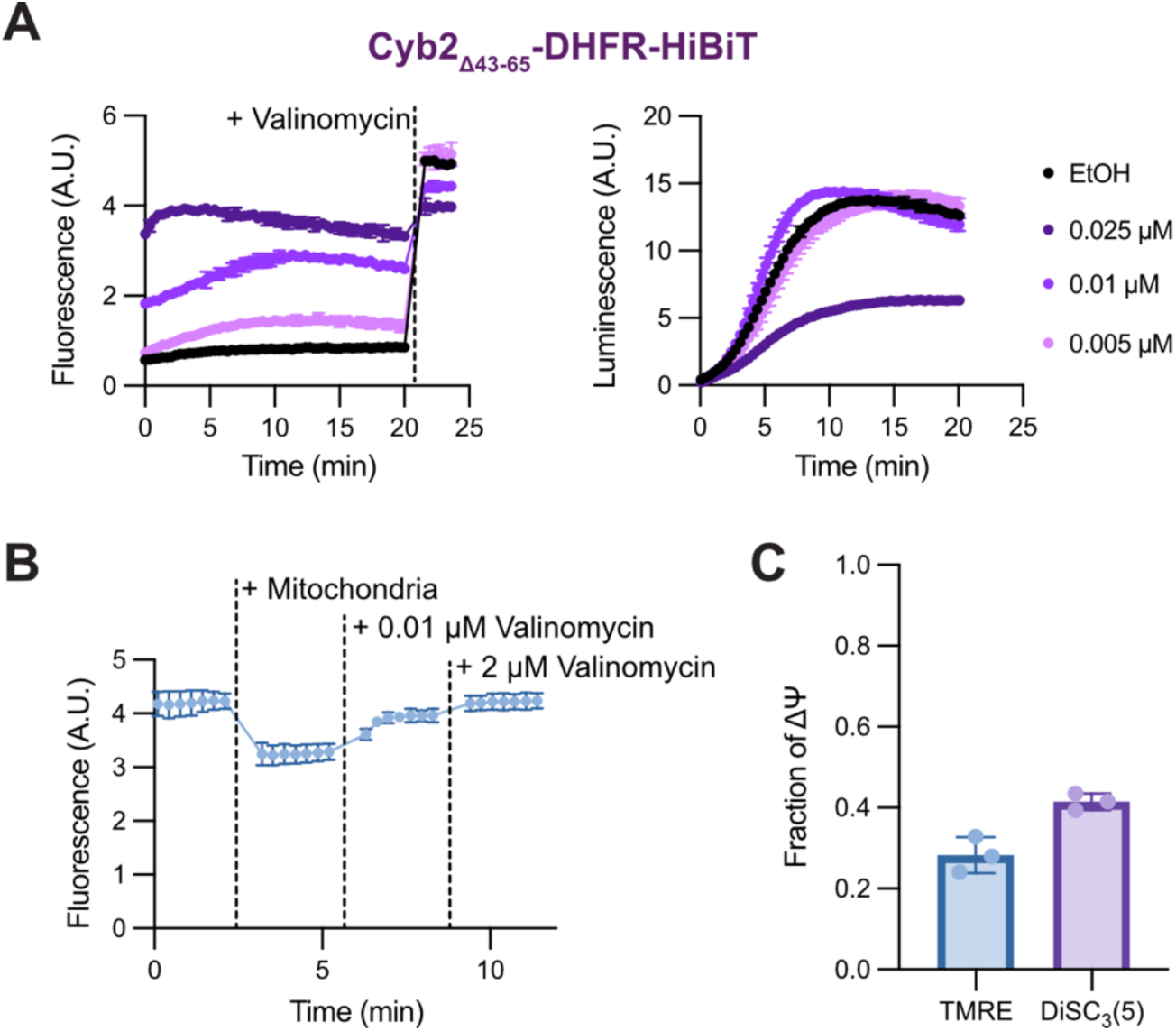
**(A)** PotLuc measurement for the import of 100 nM Cyb2_Δ43-65_-DHFR-HiBiT at varied valinomycin concentration. n = 3 independent measurements. Data represented as mean ± standard deviation. **(B)** TMRE fluorescence measurement for the effect of 0.01 μM valinomycin on mitochondrial membrane potential. n = 3 independent measurements. Data represented as mean ± standard deviation. **(C)** Quantification of data in (B) in comparison with the results of DiSC_3_(5) as in Figure 3A. n = 3 independent measurements. Data represented as mean ± standard deviation.

**Supplemental Figure 4.**
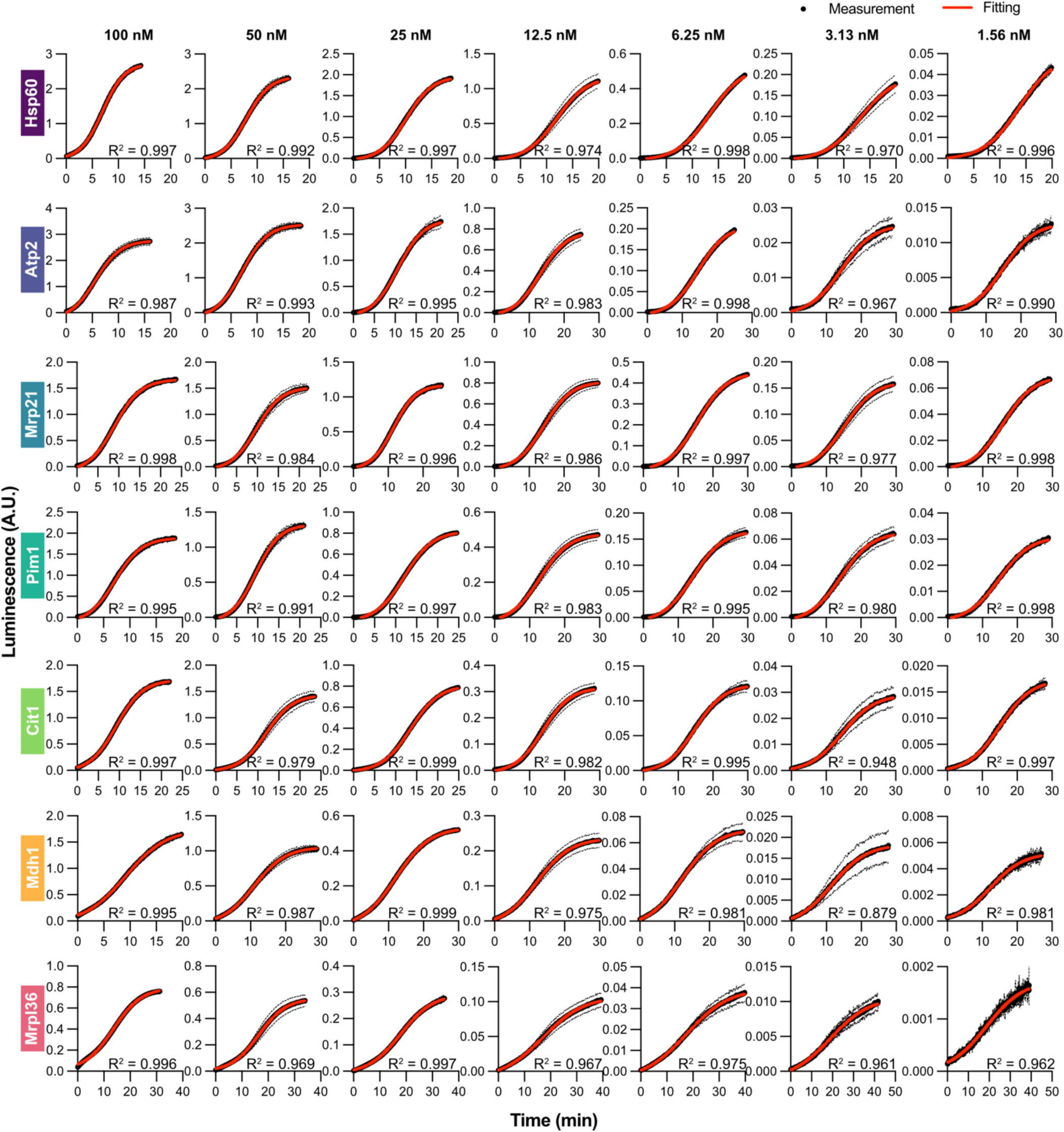
Import traces of the [presequence]-DHFR-HiBiT fusion proteins with the seven *S. cerevisiae* presequences from the MitoLuc assay were fitted with the generalized logistic function at each precursor concentration. n = 3 independent measurements. Data represented as mean ± standard deviation.

**Supplemental Figure 5.**
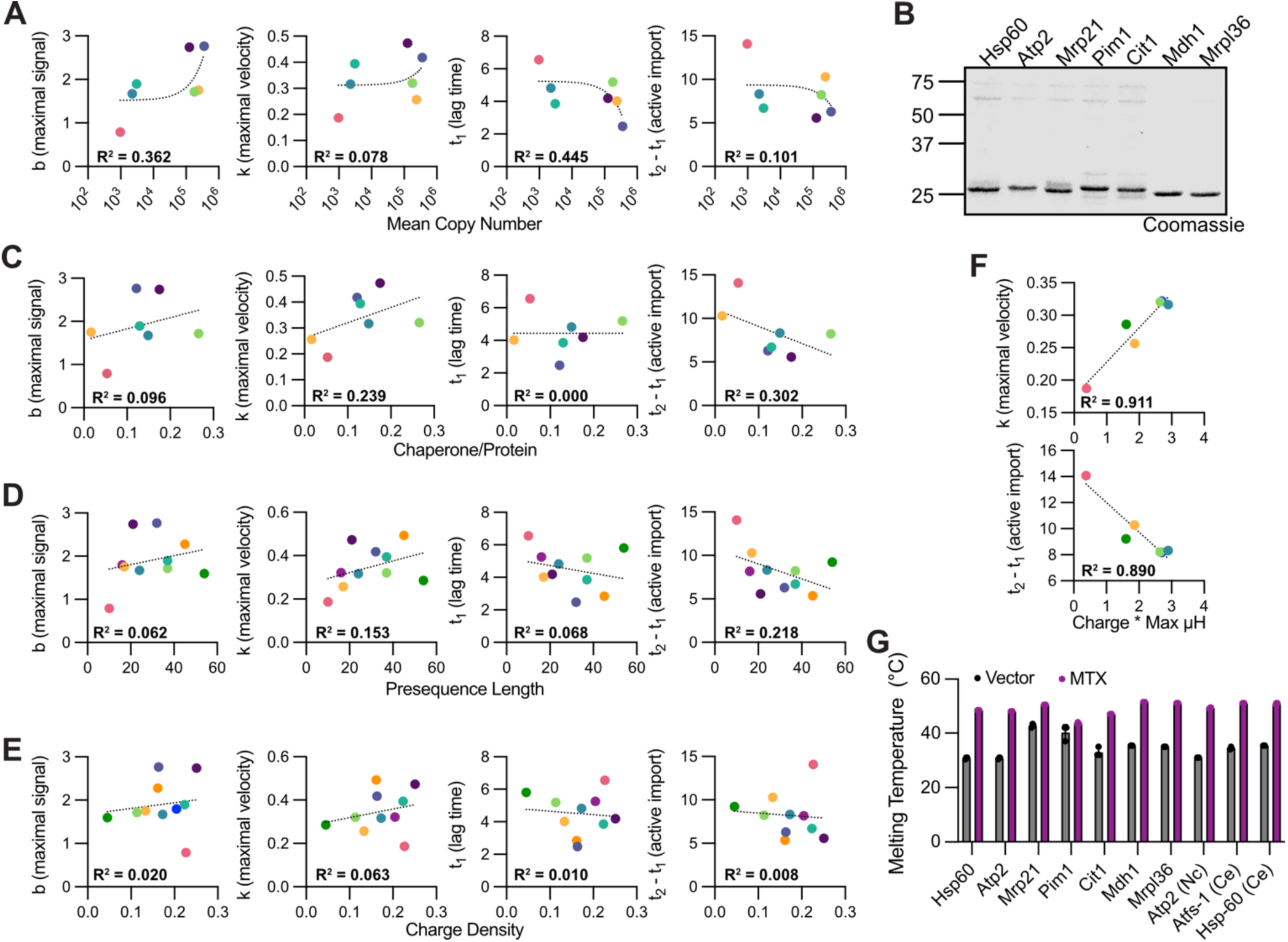
**(A)** Correlation between maximal signal (*b*), maximal velocity (*k*), lag time (*t*_1_), or duration of active import (*t*_2_-*t*_1_) at 100 nM precursor concentration and copy number of the proteins when grown on glycerol^1^. Data were fitted with simple linear regression. **(B)** SDS-PAGE for the seven precursor proteins harboring presequences from *S. cerevisiae*. Total proteins were stained with Coomassie. **(C)** Correlation between maximal signal (*b*), maximal velocity (*k*), lag time (*t*_1_), or duration of active import (*t*_2_-*t*_1_) at 100 nM precursor concentration and relative amount of chaperones based on quantification of (B). **(D-E)** Correlation between maximal signal (*b*), maximal velocity (*k*), lag time (*t*_1_), or duration of active import (*t*_2_-*t*_1_) at 100 nM precursor concentration and presequence length (D) or presequence charge density at pH 7.4 (E). Data were fitted with simple linear regression. **(F)** Correlation between maximal velocity (*k*) or duration of active import (*t*_2_-*t*_1_) at 100 nM precursor concentration and the product of maximal helical hydrophobic moment and net charge at pH 7.4 for the six proteins with lower maximal velocity (HSP-60-, Cit1-, Mrp21-, ATFS-1-, Mdh1-, and Mrpl36-DHFR-HiBiT). **(G)** Melting temperatures of the ten precursor proteins in the presence and absence of methotrexate. n = 3 independent measurements. Data represented as mean ± standard deviation.

**Supplemental Figure 6.**
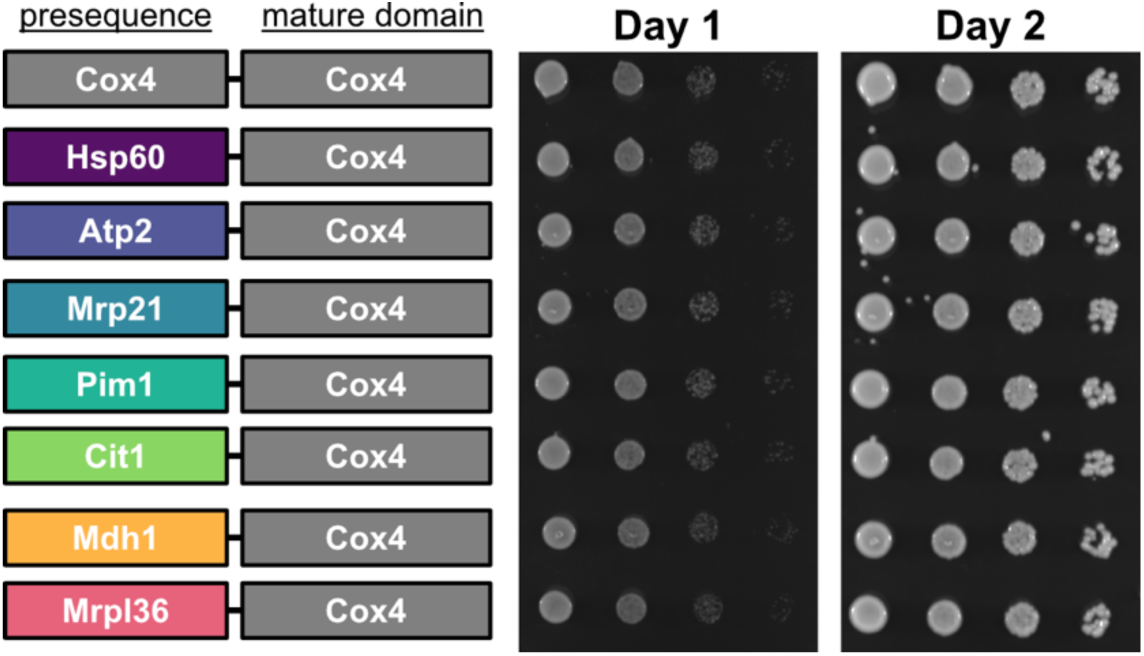
Spotting assay for comparing the growth rates of *ΔCOX4* cells rescued with Cox4 with its endogenous presequence or with the seven *S. cerevisiae* presequences tested. Serial dilutions were plated on Ura^-^ media with 2% glucose and incubated at 30°C for the indicated number of days.

